# Finding Druggable Sites in Proteins using TACTICS

**DOI:** 10.1101/2021.02.21.432120

**Authors:** Daniel J. Evans, Remy A. Yovanno, Sanim Rahman, David W. Cao, Morgan Q. Beckett, Milan H. Patel, Afif F. Bandak, Albert Y. Lau

## Abstract

Structure-based drug discovery efforts require knowledge of where drug-binding sites are located on target proteins. To address the challenge of finding druggable sites, we developed a machine-learning algorithm called TACTICS (Trajectory-based Analysis of Conformations To Identify Cryptic Sites), which uses an ensemble of molecular structures (such as molecular dynamics simulation data) as input. First, TACTICS uses k-means clustering to select a small number of conformations that represent the overall conformational heterogeneity of the data. Then, TACTICS uses a random forest model to identify potentially bindable residues in each selected conformation, based on protein motion and geometry. Lastly, residues in possible binding pockets are scored using fragment docking. As proof-of-principle, TACTICS was applied to the analysis of simulations of the SARS-CoV-2 main protease and methyltransferase and the *Yersinia pestis* aryl carrier protein. Our approach recapitulates known small-molecule binding sites and predicts the locations of sites not previously observed in experimentally determined structures. The TACTICS code is available at https://github.com/Albert-Lau-Lab/tactics_protein_analysis.

## Introduction

Designing allosteric modulators requires a knowledge of which parts of the protein can bind ligands with high affinity. This is not a trivial task, especially in cases where the binding site can be classified as “cryptic”. A cryptic site can be broadly defined as a site that is apparent when a ligand is bound to it but difficult to detect when no ligand is present(1). One proposed mechanism for cryptic site formation is the “induced fit” mechanism, in which the ligand causes the protein to adopt a conformation that it would otherwise not adopt. Another proposed mechanism is “conformational selection”, in which ligands bind and stabilize pocket-containing conformations that are rare (but not nonexistent) in unbound proteins. It is likely that many cryptic sites undergo a combination of these mechanisms(2),(3),(4). The exact definition of a cryptic pocket is subject to debate; a particularly difficult question involves the classification of sites that are visible in some unbound structures but are hidden in other structures. Some classify these sites as cryptic(5), but this is not universally accepted(1). Finding a cryptic site on a protein does not guarantee that the protein is druggable. For a cryptic site to be of interest for drug design, it must also be allosteric: the binding site’s conformation must influence the protein’s function(6). Despite this caveat, the ability to find cryptic pockets opens up an additional promising avenue for drug design.

A variety of algorithms have been developed to study binding sites (both cryptic and non-cryptic). Some algorithms characterize the behavior of known pockets. Many of these algorithms analyze molecular dynamics (MD) simulations, including EPOCK(7), POVME(8),(9),(10), and TRAPP(11),(12),(13). These algorithms are useful for characterizing the behavior of known pockets, but their reliance on prior knowledge of binding pocket locations limits their use on poorly understood proteins.

Other algorithms predict the locations of previously unknown pockets. Some take a static structure (such as a PDB file) as input, such as LIGSITE(14), KVFinder(15), COACH(16), COFACTOR(17), ConCavity(18), FPocket(19), GalaxySite (20), ghecom(21), DoGSite(22), and FTMAP(23). This approach can sometimes provide druggability scores of predicted pockets, as seen in PRANK(24) and DoGSiteScorer(25). Using a single structure reduces computational time but makes it difficult to find cryptic sites that are not present in all conformations. Using these programs on MD trajectories would require users to either run them on each frame (which is computationally expensive) or select individual frames to be analyzed (which increases the time and expertise required). Other algorithms use MD simulations; these algorithms consider multiple protein conformations and thus are sometimes able to find cryptic sites. One variety of MD procedure is “mixed-solvent MD”, in which proteins are simulated in a system containing both water and small-molecule probes(26),(27),(28),(29),(30),(31). The probes may interact with the binding sites on the protein, revealing the sites’ locations. Probes can open pockets that operate by the induced fit mechanism. The SWISH procedure(2),(32) combines the use of probes with enhanced sampling. A procedure somewhat similar to mixed-solvent MD “wraps” the protein in a coating of ligand before MD simulation(33). While these procedures are extremely powerful, they require specialized simulations. This prevents their use on precomputed trajectories.

Other algorithms that find pockets or binding sites take the results of precomputed MD simulations that may not include probes. EPOS^BP^ (34), trj_cavity(35), MDPocket(36), D3Pockets(37), and NanoShaper(38),(39) begin with geometry-based pocket finders that neglect physicochemical properties; then they analyze the newly found pockets. Pocketron(40) begins with NanoShaper, then performs detailed analyses with an emphasis on characterizing allosteric regulation. FTDyn(41) docks small probes onto frames from precomputed MD trajectories in order to find bindable regions. The exposon procedure(42) uses correlated changes in solvent-accessible surface area to find cryptic pockets. Each of these procedures can be quite useful, but all have limitations. Some do not consider physicochemical properties (such as those examined in fragment docking), leaving users uncertain about the extent to which geometrically reasonable pockets will bind ligands. Other programs run on each frame of the trajectory, making it time consuming to run the software and interpret the output for long simulations. Users can mitigate this by selecting individual frames for analysis (e.g., through clustering), but this increases the workflow’s complexity. By incorporating the abilities to both select individual frames for analysis and consider physicochemsitry into a single software package, TACTICS fulfils a unique niche.

CryptoSite(5) takes a different approach by calculating its own MD trajectories without probes within the algorithm. The JEDI procedure(43) also includes probe-free MD simulations, with the JEDI score used as a collective variable to enhance sampling. However, the procedure requires users to define a region of interest, limiting the applicability to proteins lacking any information about cryptic site locations.

Each of these strategies has advantages and disadvantages. Mixed-solvent MD probes can open cryptic pockets that operate by the induced-fit mechanism. However, mixed-solvent MD simulations must avoid undesirable interactions between probe molecules. Strategies for this include adding repulsion between probes(30), choosing probes that do not aggregate(26), and simulating each probe type separately(27). MD simulations without probes avoid the issue of probe-probe interactions, but they risk missing cryptic sites that require induced fit. Because many cryptic sites’ opening mechanisms include conformational selection (in addition to induced fit), it is possible that probe-free simulations will expose part (if not all) of the existing cryptic sites.

Developers of pocket-finding software must also consider whether their approach should analyze existing MD simulations or run MD within the algorithm. The latter strategy allows software users who are unfamiliar with MD to explore conformational space without the effort of learning to set up their own simulation. However, automatic system generation may prevent users from studying proteins that require specialized or nonstandard conditions. Software that takes precomputed MD trajectories assumes that users have the knowledge necessary to set up and run an MD simulation. However, it gives users more flexibility in the types of simulations that can be studied. Software that analyzes precomputed trajectories also permits trajectories calculated for other studies to be repurposed as TACTICS input for pocket-finding research.

The ability of machine learning (ML) to find trends in complex data makes it a promising technique for the nontrivial task of predicting small molecule binding sites. A few algorithms that find binding sites use machine learning(16),(5), but most do not. Machine learning has the potential to substantially improve binding site prediction, including prediction of “cryptic pockets” that are not present in crystal structures(1). Thus efforts to apply ML and other cutting-edge computational techniques to the prediction of binding sites can have substantial impacts on the process of designing drugs.

We designed and implemented a machine learning algorithm that finds druggable pockets within MD trajectories. The algorithm is named TACTICS (“Trajectory-based Analysis of Conformations To Identify Cryptic Sites”). As described below, TACTICS correctly identifies several known or predicted ligand binding sites in SARS-CoV-2 proteins. It also predicts several novel binding sites in the SARS-CoV-2 2’-O RNA methyltransferase and the *Yersinia pestis* aryl carrier protein. These predicted sites provide new opportunities for drug development. The TACTICS software is freely available so that it can be used to find druggable sites in other proteins.

## Methods

### ML Training Database Generation

The ML model used in TACTICS was trained using a database of crystal structures. Each protein in the database has a single “holo” structure with a ligand bound in the cryptic site and up to 50 “apo” structures without a ligand in the site. These ensembles of structures are designed to imitate MD trajectories in which a few frames have the cryptic site that is absent in most frames.

Database construction began with a reconstruction of the Cryptosite Database(5). Each cryptic site in this database has an “apo” structure with the site hidden and a “holo” structure with a ligand in the cryptic site. The original database contains the AMPAR LBD twice. Both entries were removed from the reconstructed database.

After the Cryptosite Database was reconstructed, it was extended using a procedure similar to that used in Beglov et al.(3). For each database protein, the PDB’s 95% sequence identity cluster was downloaded. This provided additional apo structures to be compared with each holo structure. This is important because the database is designed to mimic MD trajectories in which many frames may have the pocket region in conformations different from the bound conformation. While the percentage of time that a pocket is open in an MD trajectory is likely to be system-dependent, it is reasonable to suppose that ensembles of conformations generated by MD would frequently have high ratios of apo-like structures to holo-like structures. The ML training database was designed to mimic this.

MODELLER(44) was used to match the holo structure’s residue numbers with those of each structure in the cluster. PyMOL(45) was used to select residues in the holo structure within 9 Å of the co-crystallized ligand; these residues were used to align the holo structure with each extended-database structure. All extended-site structures with a ligand other than water or a metal within 5°A of the holo ligand were removed from the database. This was done because no apo structures should have the cryptic site bound.

Our procedure assumes that in the Cryptosite database, cryptic sites are closed (or otherwise unbindable) in all unbound (“apo”) structures. However, some “apo” structures may have the cryptic site partially or fully open and bindable despite the absence of ligand(3). This shortcoming of the present database may impact the ML model’s accuracy. This is because the ML training procedure seeks to identify how protein conformations differ when cryptic sites are bound or unbound; when the bound and unbound structures resemble each other, this is impossible.

Eliminating this shortcoming would be difficult: proteins exist in a spectrum of conformations with more or less accessible cryptic sites, and classifying intermediate states as either bindable or unbindable would require arbitrary decisions.

An additional shortcoming is presented by the variability in the number of apo structures per holo structure. Some proteins have been crystallized more often than others, resulting in a biased distribution of crystal structures. To reduce the severity of this issue, a maximum of 50 apo structures per holo structure was used. This does not completely eliminate the issue, as proteins with fewer than 50 structures will be undersampled. However, it reduces the issue’s severity.

### ML Model Construction

A machine learning model was constructed to find cryptic sites. The model predicts the likelihood that a given residue in a given structure is part of an accessible cryptic site. The model was designed to return False for residues that are not part of a cryptic site, and for residues that are part of a cryptic site that is not accessible in the input structure. It is only intended to return True when the input structure contains an accessible cryptic site.

The model uses several features. One feature is the ConCavity score for each residue(18). ConCavity is a pocket detection algorithm that considers the geometry of the input structure and (if allowed by the user) the sequence conservation of each residue. It has been used in other multi-step pocket finding algorithms(5),(16).

ConCavity has the option of only using the structural information (ignoring sequence conservation); TACTICS uses this option. Including sequence conservation in ConCavity may improve TACTICS’s results; this might be added to future versions of the software. ConCavity’s structure-based approach starts by using Ligsite(14) to determine which points near the protein are part of geometric pockets. ConCavity then clusters these points into binding sites and uses these sites to assign scores to each residue. It considers only a single structure.

Another feature in TACTICS’s ML model is the change in ConCavity score between the current structure and a reference structure. This feature can only be calculated from conformational ensembles such as MD trajectories, so it sets TACTICS apart from algorithms that work on a single structure. The feature was included to increase TACTICS’s ability to find cryptic sites. It is important because flexibility has been observed to be associated with cryptic site locations(3). The reference structure must be provided by the software user. It is expected that the reference structure will usually be the first frame of the MD trajectory, but another structure can be chosen (as long as it is aligned to the MD trajectory and has identical residue numbering).

A druggable cryptic pocket should include more than one residue. The model considers this by including as a feature the average ConCavity score of the residues to either side (in the sequence) of the residue in question. It might be useful to consider spatially close residues that may not be close on the sequence; future versions of TACTICS may add this capability. The model also considers the change in sequence neighbors’ ConCavity scores between the current structure and the reference structure.

The model includes the change in α-carbon position between the current structure and the reference structure. It also includes the neighbors’ average change in α-carbon position. The values of these features were calculated for each residue of each structure in the database.

The input trajectory must be aligned so that the protein’s center of mass does not move over the course of the simulation. If this is not done, then the entire protein’s motion will incorrectly be included when calculating changes in α-carbon positions. Alignment must be done by the user prior to running TACTICS.

20% of the data was randomly selected and set aside as a test set for evaluating the model’s performance. Testing data was not used to train the model. Model performance was carried out using the area under the receiver operating characteristic (abbreviated AUROC or ROC-AUC). The statistics behind the AUROC merit a brief discussion. As the confidence threshold required for the model to classify an item as True is varied, the number of true and false positives also varies. The receiver operating characteristic (ROC) is a graph of true positive rate vs. false positive rate as the confidence threshold is varied. The area under the ROC curve (AUROC) is the fraction of the time that the model will assign higher confidence to a randomly chosen positive sample than to a randomly chosen negative sample(46). The advantage of the AUROC over the (conceptually simpler) percent accuracy metric is that the latter is biased if the dataset has unequal numbers of positive and negative samples. The AUROC has been used to quantify the performance of other pocket-finding algorithms(5),(16); it was selected to quantify the performance of TACTICS’s ML model.

Scikit-learn(47) was used to train a random forest model on the 80% of the data that was not set aside. Three-fold cross-validation was used to select hyperparameter values. After training the model on the training dataset, the AUROC score was calculated on the test dataset. A result of 0.856 was obtained. This AUROC was calculated on a dataset of crystal structures, while the ultimate application of the model is to MD trajectories. Our use of this score assumes that the conformations in the crystal structure dataset are representative of what would be seen in MD simulation data. The extent to which this assumption is true impacts the extent to which our AUROC can be assumed to represent performance on MD data. Nevertheless, the score is useful as a rough estimate of whether the model is likely to be useful. After calculating the AUROC, the model was re-trained on the entire database (composed of both the training and testing sets).

### Incorporating The ML Model Into A Larger Algorithm

As useful as the ML model is, it was observed to predict an unrealistic number pf pockets. While it is difficult to quantify the number of false positives without a complete knowledge of which parts of the protein are bindable, models that predict overly large numbers of pockets are likely to be flawed(3). To mitigate the issue of falspe positives, the ML model was incorporated into a more complex algorithm (TACTICS). The algorithm starts by using MDAnalysis(48),(49) to perform k-means clustering on the MD trajectory. Then the ML model is run on the centroid of each cluster. Instead of outputting a binary True/False prediction, the ML model is used to predict the probability of each residue being in a cryptic site. The algorithm rejects the prediction if the confidence is below a user-specified threshold; the default value is 0.8. This high value reduces the number of false positives. The algorithm also rejects the prediction if the standard deviation of ML scores among all clusters of the trajectory is below a certain threshold. The default value is 0.25. This is done because cryptic pockets are expected to open and close; a residue that always has a high score is unlikely to be part of a cryptic pocket.

The above procedure generates a set of residues that are predicted to be part of cryptic pockets. These residues are clustered based on their proximity to each other; the result is a set of predicted binding regions. To investigate which predicted sites are likely to be druggable, fragment docking is performed on each site. AutoDock Vina(50) is used for the docking; the fragments are the 16 molecules used by FTMap(41) and Cryptosite(5): acetaldehyde, acetamide, acetone, acetonitrile, benzaldehyde, benzene, cyclohexane, dimethyl ether, ethane, ethanol, isobutanol, isopropanol, methylamine, phenol, urea, and N,N dimethylformamide. Docking is performed in a box surrounding the predicted binding site; the default is to add 8 Å in each dimension. As in Cryptosite, each residue’s “dock score” is calculated by counting the number of times that a docked fragment ligand is within 3.5 Å of the residue. The dock score is used here to verify and expand upon the ML predictions. It is expected that predictions are more likely to be accurate when the residues with high ML scores are in the same region as residues with high dock scores. Substantial differences between docking and ML results may indicate inaccurate ML predictions. They may also represent cases where docking has found additional residues that are important for binding but were missed by the ML model. Figure 1A shows a flowchart of the TACTICS algorithm.

**Figure 1.**
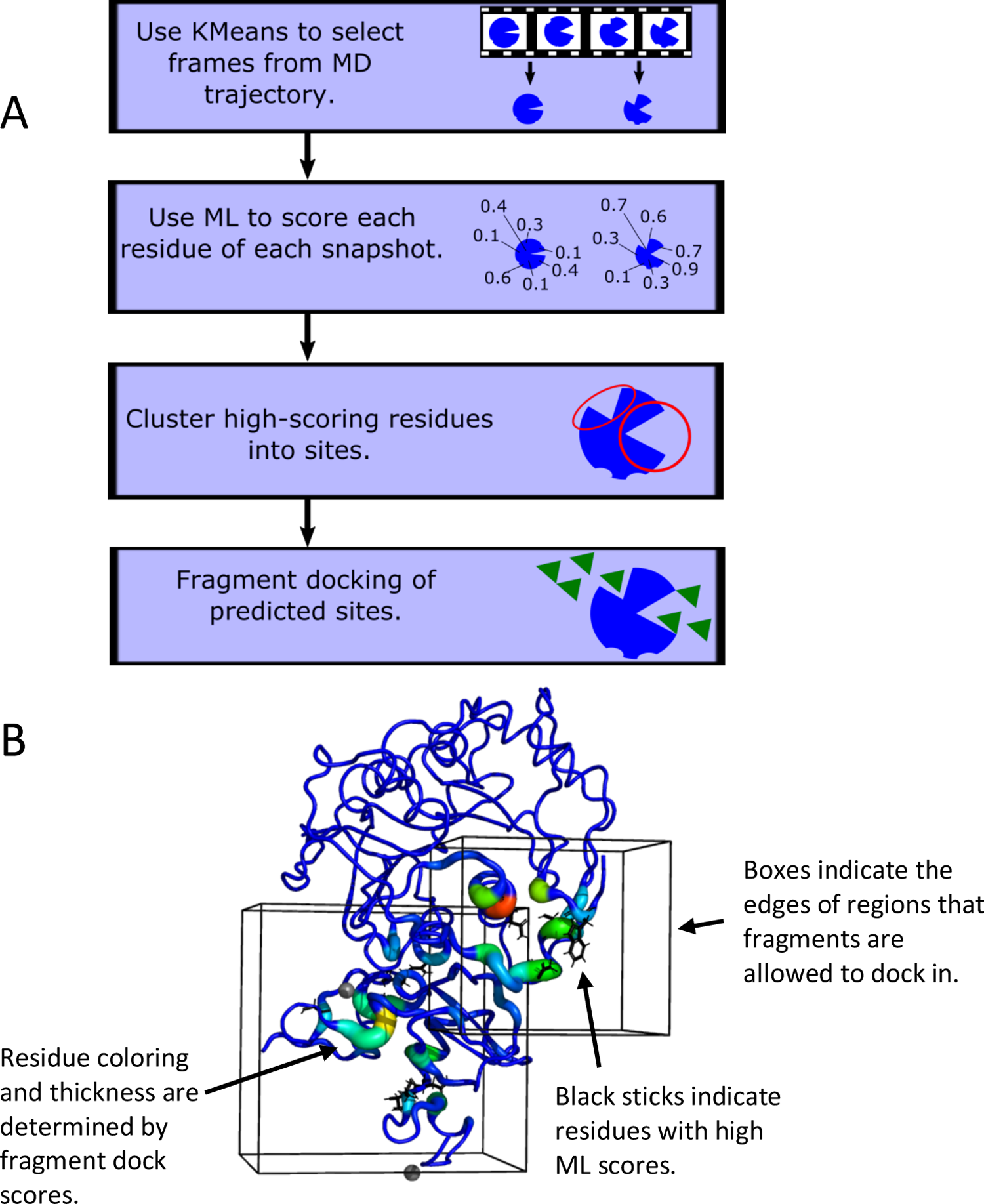
The design of TACTICS. A. Flowchart describing the TACTICS algorithm. B. Sample image from typical TACTICS output, with labels emphasizing key features.

### Visualizing the Results

To facilitate the process of interpreting TACTICS’s results, a procedure was developed for visualizing the results in PyMOL(45). Residues that are highlighted by the ML algorithm are shown as sticks. Dock scores are shown as b-factor putty. The “search spaces” where docking ligands are allowed to be are drawn as boxes. Each time that the code runs, it creates a file with PyMOL commands to display the results. Figure 1B shows typical TACTICS output.

### SARS-CoV-2 Simulation System Preparation

The SARS-CoV-2 main protease (nsp5) model is based on PDB ID 6Y2E(51). Hydrogens were added using CHARMM(52). The system was neutralized and solvated with ∼150 mM KCl for a total system size of 126,736 atoms. A weak center of mass restraint was applied to N, CA, and C atoms of residues 19-21, 36-38, and 80-82 during equilibration. No restraints were used during production simulations.

The initial atomic model of the SARS-CoV-2 2’-O RNA methyltransferase (nsp10/nsp16 heterodimer) is based on PDB ID 6W61. It was prepared in CHARMM-GUI(53). Missing amino acids in nsp10 were added using the Modloop server(54). All crystallographic waters and coordinated Zn^2+^ ions were included. The system was neutralized and solvated with ∼150 mM KCl for a total system size of 65,958 atoms. Zn^2+^ coordinating cysteines were deprotonated but retain a neutral charge using a custom force field topology. This was done to mimic the geometry of the coordination bond without introducing a nonexistent negative charge on the sulfur atoms. To prevent the dislodging of the Zn^2+^ ion from its binding cavity, a 10 kcal/mol/Å^2^ harmonic restraint was applied between the Zn^2+^ ion and each coordinating cysteine residue. A 1 kcal/mol/Å^2^ harmonic potential position restraint is applied to Cα atoms in core regions of the nsp10/nsp16 dimer (residues 70-72, 86-88, and 171-173 in nsp16) to keep the protein centered within the box. These restraints were used during both equilibration and production.

### SARS-CoV-2 REST-2 Simulations

All REST2 simulations(55) were carried out in NAMD2.13(56). The CHARMM36 force field was used for protein and ions(57),(58),(59) while the TIP3P model was used for water(60). The nsp10/nsp16 dimer and the protease were subjected to energy minimization and a 1 ns equilibration under NPT conditions at 1 atm and 300 K. All REST2 simulations used an NPT ensemble at 1 atm, with the effective temperature ranging from 300 to 350 K, spanning across 12 replicas for each system. Solute tempering is applied to protein atoms only and exchange attempt frequency is set to every 2 ps. A 1 ns pre-production REST2 simulation was carried out to initialize exchange probabilities. Each replica was simulated for 50 ns to obtain exchange probabilities between 20% and 30%.

For equilibration, pre-production, and production simulations, a timestep of 2 fs was used. Periodic boundary conditions were used. All electrostatic interactions were computed using the particle-mesh Ewald (PME) algorithm, and short-range, non-bonded interactions were truncated at 12 Å.

### ArCP Simulations

The Aryl Carrier Protein (ArCP) simulation systems are based on PDB structures 5TTB (apo)(61), 2N6Y (holo)(62), and 2N6Z (loaded)(62). The loaded and holo structures each have a phosphopantetheine (PP) “arm” attached to Ser52 and the loaded form harbors a salicylate at the end of this arm; these moieties were parameterized using CGenFF(63),(64),(65). Each protein (apo, holo, and loaded) was minimized in a vacuum before being solvated in a water box whose sides are 60Å wide. 17 sodium atoms and 18 chlorine atoms were added, neutralizing the system and providing a concentration of 150 mM NaCl. Each system underwent three rounds of 60,000 ps equilibration at 300K using Langevin dynamics. Equilibration and production simulations used an NPT ensemble.

For each system, the simulation was run in NAMD using the CHARMM36 force field. The simulations were carried out at 300 K and 1 atm with periodic boundary conditions. A timestep of 2 femtoseconds was used, and a trajectory snapshot was written every 500 timesteps (1 ps). A total of 200 ns of simulation data was obtained, and the latter 100 ns was used for TACTICS analysis.

### Running TACTICS

While default values exist for TACTICS parameters, these values may not be optimal for all systems. Poor parameter choices may lead to TACTICS not finding few (if any) pockets or finding unreasonably many pockets. Finding the best parameter values may require running TACTICS with several values and evaluating which run has the most reasonable results. This introduces an unfortunate subjectivity into the TACTICS workflow. But eliminating all user-defined parameters would risk compromising the software’s ability to generate useful results across a diverse range of input protein systems.

The tunable parameters are as follows. `num_clusters` is the number of frames from the trajectory that will be selected for TACTICS analysis. Frame selection uses k-means clustering. The number of frames displayed in the TACTICS output may sometimes be less than `num_clusters`; this occurs if there are frames with no pockets. Values of roughly 3-15 are expected to be reasonable for most situations; it is recommended to start with a value near 8. `ml_score_thresh` is the minimum ML score needed to be considered when clustering residues into binding sites. The default value is 0.8; substantial variation may be needed but values of 0.5-0.9 should work for most systems. `ml_std_thresh` is the maximum standard deviation between frames that a residues ML score must have for the residue to be considered when clustering residues into binding sites. It may also benefit from substantial variation; the default value is 0.25 and values from roughly 0.05 to 0.4 are suggested.

It should be emphasized that the trajectories should be aligned before running TACTICS. This removes the motion of the entire protein that would affect the ML model’s change-in-position features. More information on how to run TACTICS can be found in the software’s README file.

For the main protease, we used num_clusters=8, ml_score_thresh=0.6, and ml_std_thresh=0.09. For the methyltransferase, we used num_clusters=8, ml_score_thresh=0.6, and ml_std_thresh=0.1875. For the ArCP, we used num_clusters=13, ml_score_thresh=0.6, and ml_std_thresh=0.15. Each trajectory was aligned using MDAnalysis(49),(48).

Using TACTICS to analyze large datasets is computationally intensive. Thus we divided the data into chunks and ran TACTICS on each chunk. For each SARS-CoV-2 protein, we ran TACTICS on the first 20 ns, the next 20 ns, and the last 10 ns. Analyzing the ArCP simulations was less computationally demanding, so these simulations were not divided into chunks.

### Analyzing Proteins With Non-Standard Components

Our SARS-CoV-2 methyltransferase simulations included 2 zinc ions. Our ArCP simulations included a covalently attached phosphopantetheine (PP) arm. These atoms can be expected to impact ligand binding to any nearby sites by, for example, steric hindrance. Ignoring these non-standard components during TACTICS analysis might introduce inaccuracies. Including them in TACTICS analysis required special consideration; the procedure is expected to be useful for other proteins with non-standard components.

TACTICS requires each residue name be one of the standard amino acids. Additionally, each residue must have an alpha carbon (atom name CA). TACTICS deletes atoms that do not meet these conditions. A relatively straightforward way of getting TACTICS to recognize coordinated zinc atoms is to change their residue names from Zn to an amino acid and change their atom names to CA. Based on how TACTICS works (including ConCavity and AutoDock Vina, which it runs), we believe that approximating coordinated zincs as alpha carbons is reasonable. Because both methyltransferase zinc atoms coordinate with multiple residues in the protein, we expect the zincs’ electrostatic effects to be minor enough that ignoring them does not dramatically impact our results. (Atoms with strong hydrophobicity or hydrogen-bond formation favorability would impact fragment docking.) The alpha-carbon approximation acknowledges that zinc atoms create excluded volume; thus, making the approximation is preferable to letting TACTICS delete the zincs.

In the case of the ArCP, the loaded and holo forms have a PP arm covalently attached to a serine residue. The modified residue already has an alpha carbon, but it must be renamed to an amino acid (e.g. Ser) before TACTICS analysis. We also had to rename an atom with atom name “CS”; the atom is supposed to be carbon but TACTICS attempted to parse the “CS” as cesium.

## Results

To demonstrate the capabilities of TACTICS, we applied the algorithm to three proteins. Two of them are SARS-CoV-2 proteins (main protease and methyltransferase). TACTICS finds previously reported binding sites and novel sites in these proteins. The third protein (aryl carrier protein) is relatively small and has significant posttranslational modifications; it illustrates the versatility of TACTICS for unusual systems. The SARS-CoV-2 proteins were simulated using REST2(55), while the aryl carrier protein was simulated using equilibrium MD. This demonstrates TACTICS’s ability to analyze conformational ensembles generated through a variety of methods.

### SARS-CoV-2 Main Protease

The SARS-CoV-2 main protease (M-pro; nsp5), is responsible for cleaving the polyprotein produced from translation of the viral RNA genome. Specifically, M-pro cleaves the peptide bond after glutamine residues most frequently occurring in the recognition sequence Leu-Gln-(Ser, Ala, Gly)(51) (66). Peptide cleavage by M-pro occurs by a catalytic dyad of cysteine and histidine residues in the active site(67).

While cleavage after glutamine residues has been observed for the main proteases of other coronaviruses, including SARS-CoV(68) and MERS(69), it has not been frequently observed in human proteases(66). From a therapeutic perspective, this reduces the probability of off-target effects and makes M-pro a promising target for drug design. Most notably, the binding of peptidomimetic inhibitors to the proteolytic active site has been explored as a means to inactivate the protease(70). By mimicking the geometry of the substrate, these inhibitors become “trapped” in the active site, often by the creation of a covalent bond with the catalytic cysteine(70), thus inactivating the protein. M-pro is an obligate homodimer stabilized by interactions between the N-terminal “finger” residues of one monomer and a glutamate residue in the active site of the opposite monomer(51) (71). The requirement of dimerization for proteolytic activity presents another opportunity for drug development(72); the dimer interface has been more recently explored as a potential allosteric drug-binding site(73).

M-Pro has been an especially common target for structural-biological efforts to find allosteric pockets. MD-based approaches have been applied to M-Pro(74),(75),(76); large-scale crystallography screens have also been carried out(77),(78). Because the protein’s ligand-binding characteristics are relatively well-studied, M-Pro is a good choice for examining TACTICS’s ability to rediscover known binding sites. To obtain an ensemble of M-pro conformers suitable for analysis with TACTICS, we performed replica exchange with solute tempering (REST2) molecular dynamics simulations of the apo M-pro dimer. This is useful for assessing TACTICS’s veracity and usefulness.

TACTICS identifies the main protease active site. At various points in the simulation, TACTICS identifies residues including H41, F140, and H163. These residues are important for binding, as shown by their interactions with inhibitors that bind to the active site (51),(79), (80) and simulated interactions with a peptide substrate(81). The binding site is shown in Figure 2 A, B and Supplemental Figure S1. To the left of the crystallized ligand in the figure, TACTICS identifies additional residues as druggable. Many of these residues were also highlighted in the Folding@Home results(74), underscoring the likelihood that these residues are bindable. Because TACTICS was designed with an emphasis on finding cryptic sites and minimizing false positives, it uses `ml_std_thresh` to filter out residues whose scores don’t change. Thus it is not obvious that TACTICS would be able to find active sites (which might be expected to remain visible throughout the simulation). However, enzymes can be highly flexible, particularly when unbound(82).

**Figure 2.**
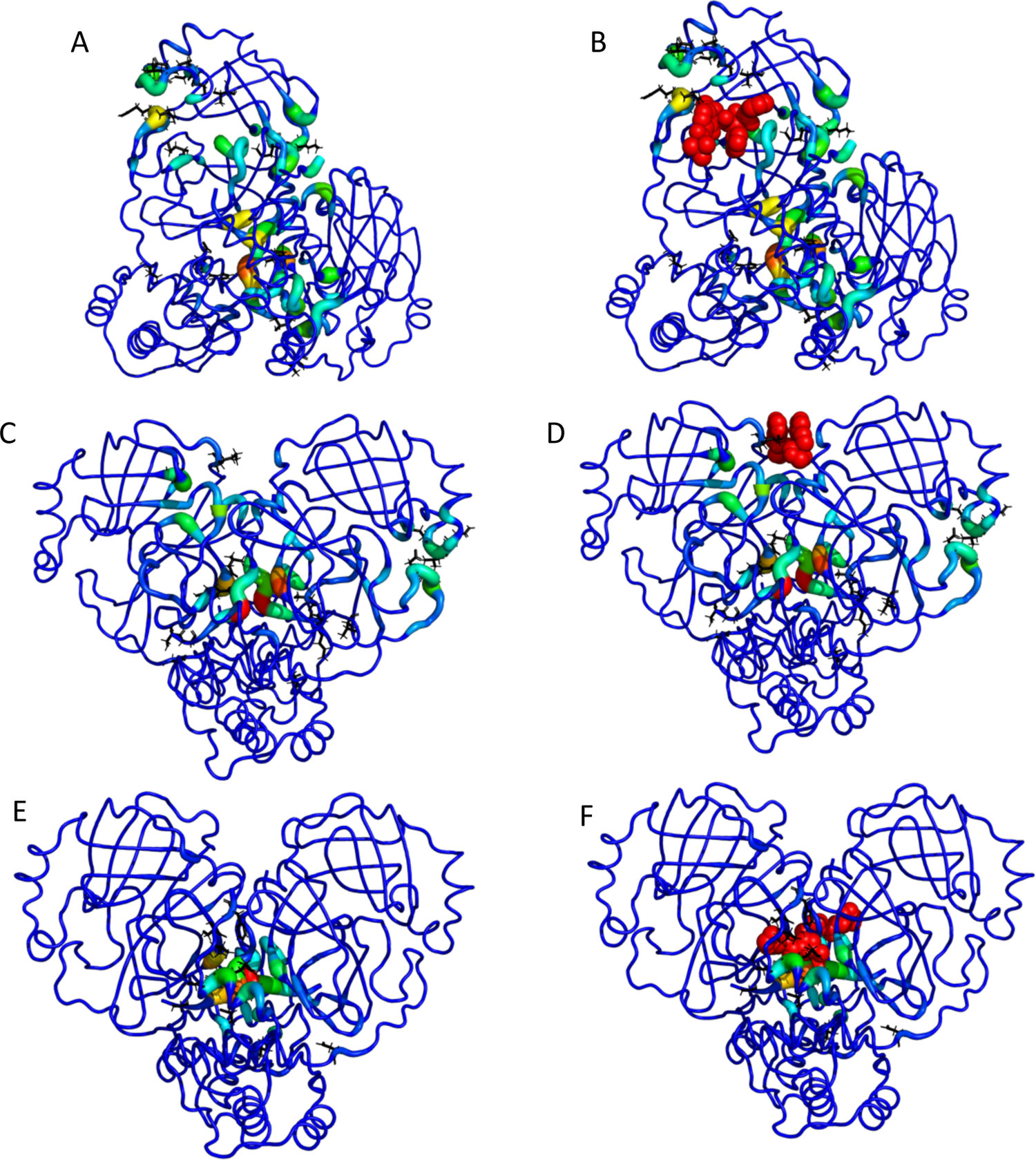
Selected images from the TACTICS output for the SARS-CoV-2 main protease. A. The TACTICS output for a frame in which the active site is found. B. The same as (A), with the active-site inhibitor called 13b added from PDB structure 6Y2F. C. TACTICS output for a frame in which a known allosteric site is found. This site binds a ligand called x0425 or Z1401276297. The x0425-binding allosteric site is found in the teal and green residues at the top of the image. In this image, the dimer interface is also high-scoring (red residues in center). D. The same as (C), with the known inhibitor called x0425 or Z1401276297 added from PDB ID 5RGJ. E. TACTICS output for a frame in which a known binding site at the dimer interface is found. F. The same as (E), with the known inhibitor x1187 or Z2643472210 added from PDB ID 5RFA.

Additionally, `ml_std_thresh` was set to a low value for the main protease. Thus TACTICS’s ability to find the active site is not entirely unexpected. This success is an example of the software’s versatility.

TACTICS also identifies a known allosteric site in the main protease. A ligand is crystallized at this site in PDB ID 5RGJ(77); while the crystal structure has a ligand bound to each protomer of the dimer, the two ligands are extremely close to each other. This ligand has been shown to decrease the protease’s activity(73). The binding site is shown in Figure 2 C, D and Supplemental Figure S2. In the conformation shown, the pocket is larger than needed to accommodate the ligand from 5RGJ. Thus it is plausible that larger ligands could bind to this site.

The dimer interface is another potential drug target identified by TACTICS. Dimerization is expected to be important to protease activity(83), so ligands that bind at the dimer interface and disrupt dimerization are likely candidates for drugs. PDB ID 5RFA has two molecules of the compound x1187 bound at the dimer interface(77). Each molecule interacts with both protomers. Experimental results suggest that x1187 inhibits dimerization and protease activity(73), confirming the importance of the dimer interface. Results on x1187 show the complexity of the dimer interface’s behavior; x1187 binds the dimer form in the crystal structure(77) but inhibits dimerization in experimental tests(73). TACTICS correctly predicts that the region where x1187 binds is druggable. This is shown in Figure 2 E, F and Supplemental Figure S3.. The MD simulation contains C-terminal residues that are missing from the crystal structure; some of these residues would sterically clash with the ligand in its crystallized pose. It is plausible that these residues could move to accommodate the ligand in an induced-fit mechanism. Despite the uncertainty regarding the conformational and ligand-binding characteristics of this site, the fact that TACTICS identifies the site as druggable demonstrates the algorithm’s ability to find experimentally validated binding sites.

The x1187-binding site is close to where AT7519 has been crystallized(78). In some frames where TACTICS identifies the x1187 site, the predicted site location also includes some residues involved in binding AT7519. TACTICS is less successful at identifying AT7519-binding residues than x1187-binding residues. AT7519 shows relatively small antiviral activity(78). The weak antiviral activity and incomplete identification by TACTICS raise the possibility that the site could be less favorable to ligand binding than other sites.

### SARS-CoV-2 2’-O RNA Methyltransferase

During the COVID-19 pandemic, one of the many promising druggable targets is the SARS-CoV-2 2’-O RNA methyltransferase (MTase). This MTase is responsible for RNA cap formation, which is critical for viral RNA invasion. The active form of the MTase exists as a heterodimer of the catalytic nsp16 and the activating zinc finger protein nsp10 (84, 85). Currently, two druggable sites, universal to all protein and nucleotide methyltransferases, have been identified: the SAM cofactor and the RNA cap substrates (84, 86). Indeed, several small-molecule inhibitors have been designed to exploit these binding sites with high efficacy in other protein and nucleotide methyltransferases (87, 88). Although little functional information exists for the SARS-CoV-2 MTase specifically, it shares a striking 94.5% sequence identity with the SARS-CoV-1 MTase (89), which has been functionally well-characterized (90). Therefore, another potential druggable site-specific on the SARS-CoV-2 MTase is the dimer interface of nsp10 and nsp16, as SARS-CoV-1 MTase activity can be fully diminished by subtle alteration in the interfacial contacts of nsp10 and nsp16 (90). Although the SARS-CoV-2 MTase crystal structure shows that the interface consists of a large network of polar and hydrophobic contacts, this interface may potentially harbor potent cryptic pockets that can be revealed through MD simulations. Here, we utilize replica exchange with solute tempering (REST2) (55), a Hamiltonian replica exchange method, to effectively sample the conformational space of the SARS-CoV-2 MTase in order to identify known and cryptic binding sites with TACTICS.

Crystal structures of the SARS-CoV-2 MTase in complex with an RNA cap and Sinefugin, a SAM cofactor analog and MTase inhibitor, have been resolved (84, 85), serving as benchmarkable sites for TACTICS. In the apo form of the MTase, both the RNA cap and SAM cofactor binding sites are already able to accommodate their respective molecule.

Some of the residues recognized by TACTICS have been previously reported to impact RNA binding. At various points in the simulation, TACTICS recognizes residues including Y132, K170, H174, and E203. All of these residues were found to impact RNA binding and enzyme activity in mutation experiments on the related MERS methyltransferase(91). The RNA binding site was found in frames throughout the simulation. Figure 3 A, B shows the TACTICS output for a frame in which the RNA site is found.

**Figure 3.**
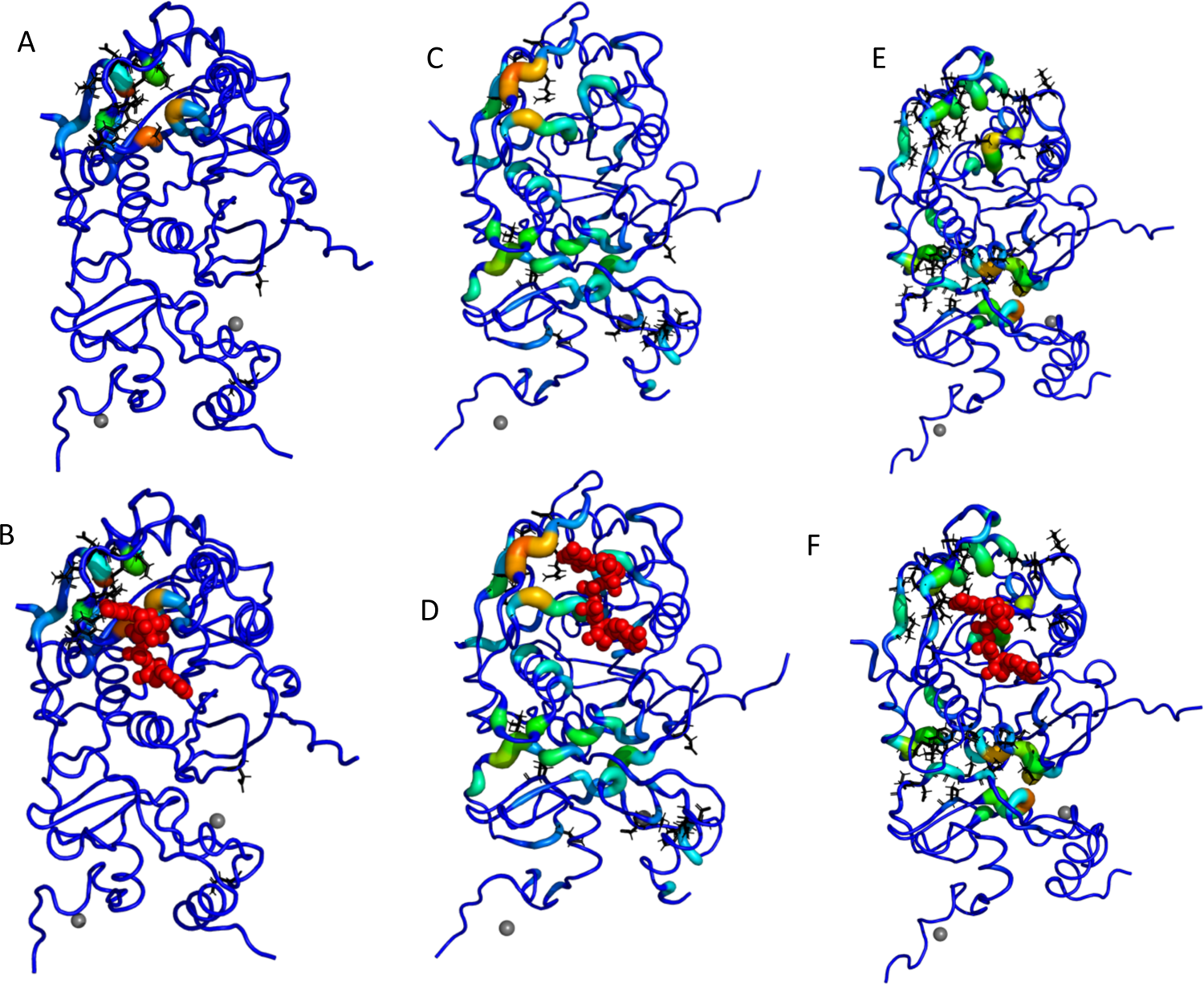
TACTICS locates the MTase RNA binding site and predicts pockets connected to it. A. TACTICS output for a frame of the MTase simulation. The RNA binding site is at the top left; it is colored by TACTICS. B. The same as (A), with the RNA Cap-0 analog m7GpppA added from PDB structure 6WVN. This confirms that the site identified by TACTICS is the RNA binding site. C. TACTICS output for another frame of the MTase simulation. D. The same as (C), with m^7^GpppA added from PDB ID 6WVN. An additional, connected site is visible to the left of m^7^GpppA. E. TACTICS output for a third frame of the MTase simulation. F. The same as (E), with m7GpppA added from PDB ID 6WVN. An additional, connected site is visible at the top right of m^7^GpppA.

TACTICS is most successful at identifying the part of the binding site that is farthest from nsp10 and the catalytic site. In addition to identifying residues known to bind RNA, TACTICS identifies conformations in which the RNA-binding pocket contains additional residues. In some conformations, certain residues in the loop between helices *α*9 and *α*10 (residues 235-241) are included in the pocket; key residues include N235, P236, and I237. The inclusion of these residues is controlled by the position of the loop between *α*2 and *α*3 (residues 17-41) and the loop between *β*8 and *β*9 (residues 196-203). L27 and S202 are particularly important for controlling access to the expanded pocket (Supplemental Figure S4F). Figure 3 C, D and Supplemental Figure S4 show this expanded pocket. In another conformation, the RNA-binding pocket includes L27, Y222, and H225; this conformation is shown in Figure 3 E, F and Supplemental Figure S5. Folding@Home(74) identifies residues near the site predicted by TACTICS; residues identified by Folding@Home include R19-D26 and K137-F152. The expanded RNA-binding sites described here include residues that do not make contact with RNA in the crystal structure. It is conceivable that a ligand binding these residues could partially fill the RNA site; it is also plausible that small ligand-induced conformational changes to the nearby region could impact the bindability of the RNA site.

TACTICS identifies the SAM-binding site, as shown in Figure 4 A, B and Supplemental Figure S6. TACTICS recognizes residues D99 and D130, both of which form hydrogen bonds with the competitive ligand Sinefungin(92). TACTICS also finds a number of nearby residues including G71, M131, and Y132. While those residues may not form hydrogen bonds (at least with sinefungin), they are still important for creating an environment that allows ligand binding. It should be noted that TACTICS finds SAM-binding residues in relatively few frames. This might demonstrate a limitation of the ability of TACTICS to find certain types of pockets. It might also suggest limitations on the number of conformations in which the SAM site is easily bindable. Further work is needed to determine which of these hypotheses are correct.

**Figure 4.**
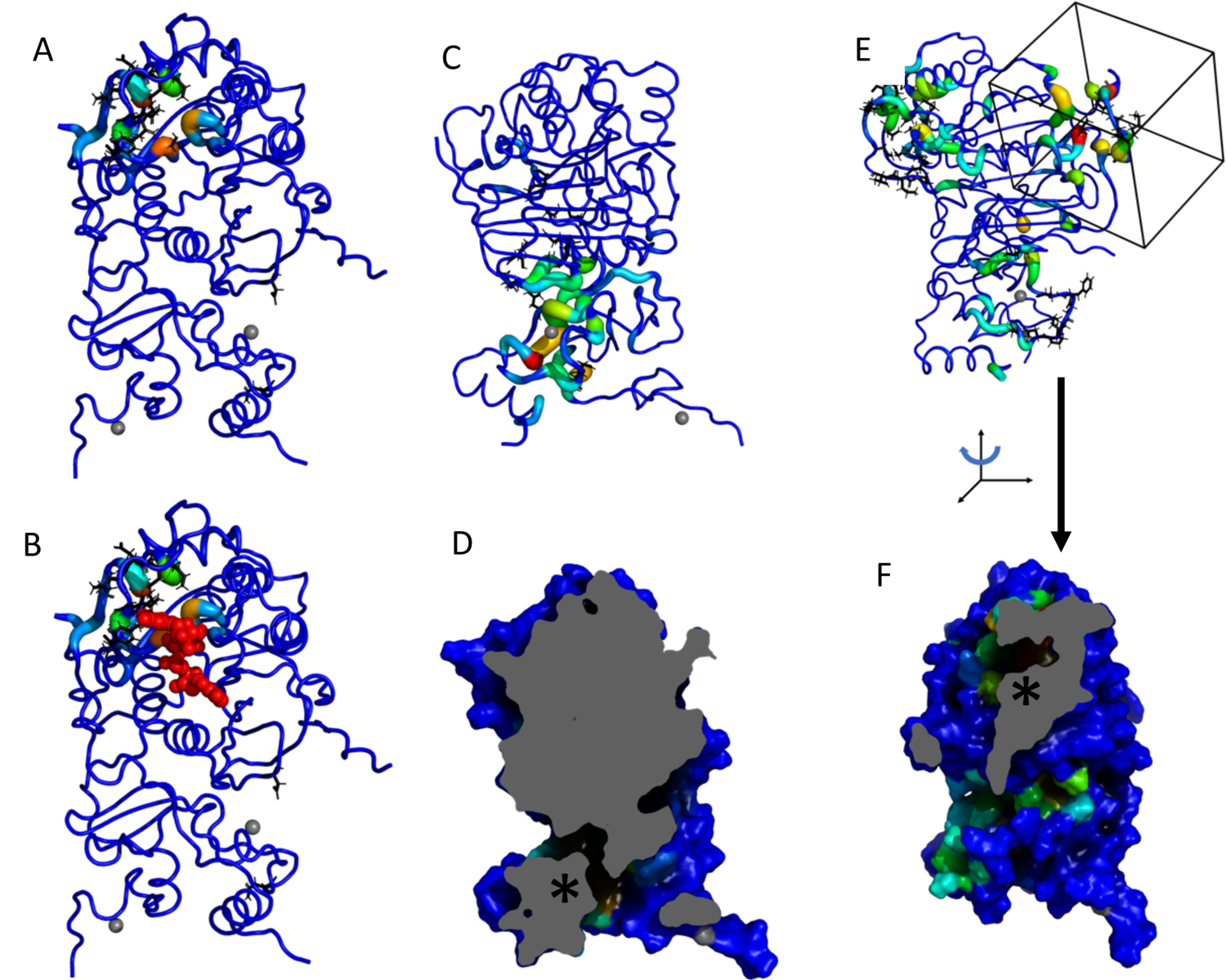
TACTICS finds additional known and predicted binding sites in MTase. A. TACTICS output for a frame in which the SAM binding site is found. The SAM binding site is at the top middle. Note that the RNA site is also colored (at the top left). B. The same as (A), with SAM added from PDB ID 6W4H. C. TACTICS output showing a predicted binding site at the dimer interface. D. Slice representation of the frame shown in (C). This image shows the tunnel through NSP10 to the dimer interface. The asterisk is next to the predicted binding site location. E. TACTICS output showing another predicted binding site. The box shows the location of the predicted binding site that has not been reported experimentally. The RNA site is colored on the left of the image. F. Rotated slice representation showing the predicted binding site. The asterisk indicates the site location.

TACTICS predicts a binding site at nsp10 and the nsp10/nsp16 dimer interface. The site is shown in Figure 4 C, D and Supplemental Figure S7. Potentially important residues on nsp10 include S72, C74, Y76, T111, L92, while potentially important residues on nsp16 include V78, S105, and D106. In some conformations (such as the one shown), the site involves a tunnel through nsp10 to the dimer interface.

In addition to the site at the dimer interface, TACTICS identified another site that may potentially disrupt activity through allosteric inhibition. The novel predicted binding site is shown in Figure 4 E, F and Supplemental Figure S8. It is on the opposite side of the protein from the RNA binding site. Potentially important residues include K123, L163, Q266, R283, and E284. A sulfate has been observed here in the crystal structure 6WVN(93), but to our knowledge no druglike ligands have been crystallized here.

### Aryl Carrier Protein

Aryl carrier protein (ArCP) is the first carrier protein domain found in the multidomain protein HMWP2, which is part of yersiniabactin synthetase, the enzymatic system producing yersiniabactin in *Yersinia pestis*.(62), (61).

Yersiniabactin synthetase is an example of a nonribosomal peptide synthetase (NRPS), which are enzymatic systems in bacteria and fungi that produce a vast number of secondary metabolites. NRPSs are composed of multiple modules organized in assembly-line fashion, such that each module is responsible for adding a single amino or aryl acid onto a forming peptide. A basic module typically contains a carrier protein (CP), an adenylation domain, and a condensation domain(62). CPs covalently tether a substrate (either the growing peptide, or the molecule being incorporated onto the peptide) via a phosphopantetheine (PP) moiety(62).

The ArCP tethers a salicylate moiety, and, being the first CP domain, can only exist in one of 3 states: apo, holo, or salicylate loaded. Apo-ArCP is inactive and must be activated via the attachment of PP to S^52^ by a phosphopantetheinyl transferase(62). Holo-ArCP refers to ArCP with the PP arm attached. The PP provides a thiol group by which salicylate can be tethered to the ArCP via a thioester bond(62). Loaded-ArCP refers to ArCP with salicylate attached to the PP arm.

Salicylate is tethered onto holo-ArCP (catalyzed by the stand-alone adenylation domain YbtE), generating loaded-ArCP(62). The cyclization domain Cy1 then catalyzes peptide bond formation between salicylate on ArCP and cysteine on the next CP domain PCP1(62) as well as subsequent cyclodehydration. In the process, salicylate is release from ArCP, regenerating holo-ArCP, and PCP1 harbors hydroxyphenylthiazoline at the end of the reaction. These steps do not occur through a rigid domain architecture; instead, during synthesis ArCP visits its partner catalytic domains through a series of transient interactions. Thus, finding small molecule binding sites on ArCP could disrupt synthesis by disrupting domain communication.

All three forms of ArCP demonstrate a four-helix bundle, with α1, α2, and α4 lying parallel to one another(61). α3 is shorter than the other helices and lies at the significant angle(61). Loop 1 connects α1 and α2. The PP binding site (Ser^52^) is located at the N-terminal end of α2(62). In both holo and loaded-ArCP, the PP arm interacts transiently with the protein core; thus, the bound state (with the PP arm docked against the protein core) exists in equilibrium with the unbound form (the PP arm not docked)(62). Thus, ArCP is a model system to assess TACTICS ability to account for such prosthetic groups. The ArCP does not have an active site, nor does it have known small-molecule binding sites (besides the PP arm). While it does have known protein binders, protein-protein binding sites have substantial differences from ligand binding sites(94).

In the apo-ArCP, TACTICS predicts two binding sites. One predicted site involves helices *α*1 and *α*2; it is shown in Figure 5 A, B and Supplemental Figure S9. Potentially bindable residues on *α*1 include Q23, R27, and E31 while key residues on *α*2 include R54, R57, and W61. As seen in the slice image, the site is divided into two distinct regions. R27 and R57 cause this division. The other predicted site is located near the N and C termini. The site is very flexible; a variety of conformations were observed. In one conformation (Figure 5 C, D and Supplemental Figure S10), the site includes *α*1 residues H17, D20, and Y21 along with *α*4 residues M87, L88, and E93. In another conformation (Figure 5 E, F and Supplemental Figure S11), the site includes *α*1 residues R16, H17, and D20 along with *α*4 residues S91, P92 and loop2 (between *α*2 and *α*3) residue Y67. In a third conformation (Figure 5 G, H and Supplemental Figure S12), the site includes *α*1 residues D14, H17, A18, and Y21 along with *α*4 residues N84, Q85, and L88.

**Figure 5.**
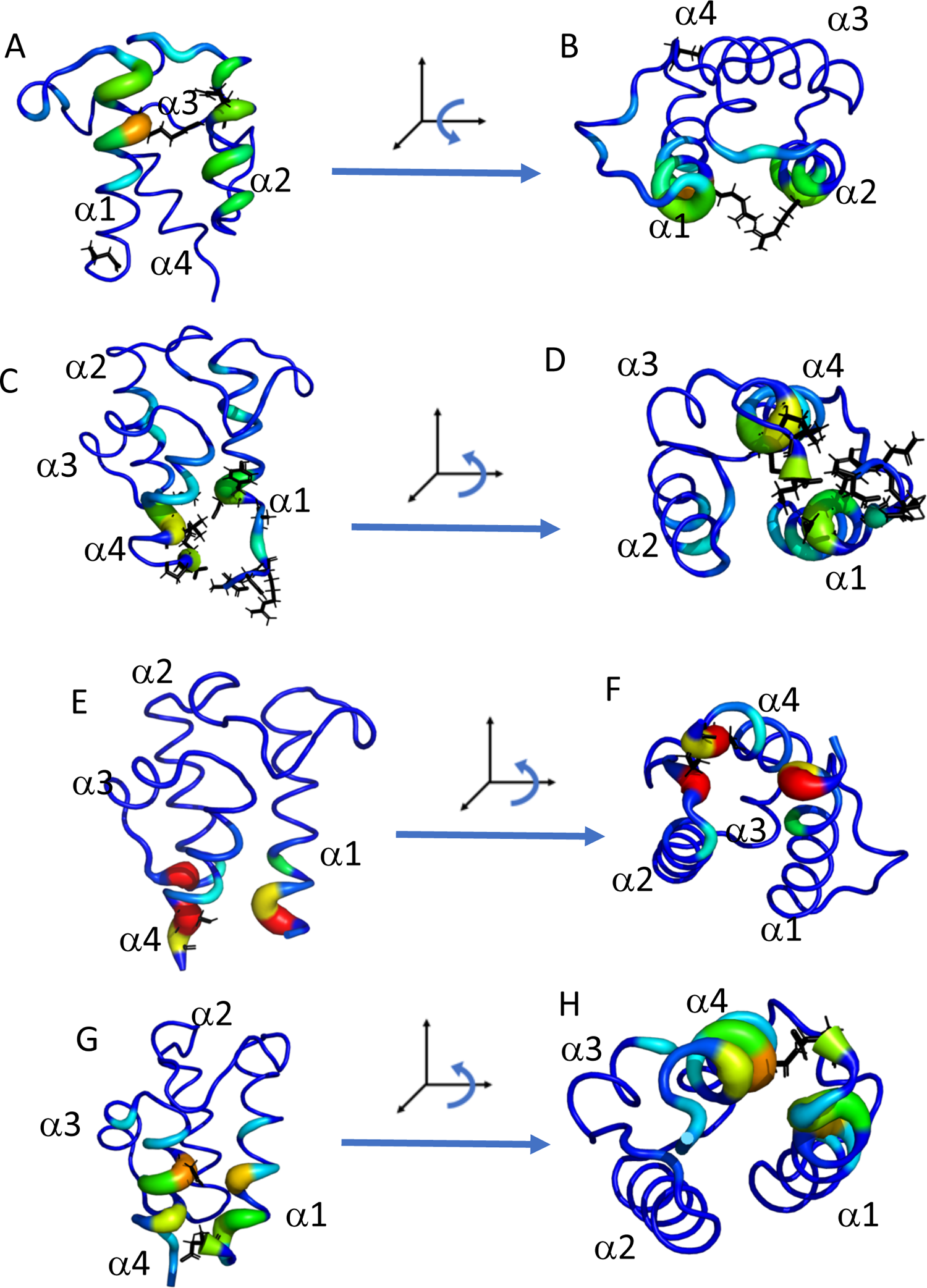
TACTICS output showing predicted binding sites in apo-ArCP. A, B. TACTICS predicts a site at the top of ArCP. C, D. TACTICS predicts a site at the bottom of ArCP. E, F. Another conformation of the site shown in (C) and (D). G, H. A third conformation of this site.

In holo-ArCP, TACTICS predicts a binding site at the N and C termini. This site is shown in Figure 6 A, B and Supplemental Figure S13. The site has the same location as the “Apo 2” site discussed above; dock scores and conformational flexibility are very similar between the apo and holo results. TACTICS also predicts a binding site located primarily on loop1 (between *α*1 and *α*2). It includes residues L39, H40, S43, and A48.

**Figure 6.**
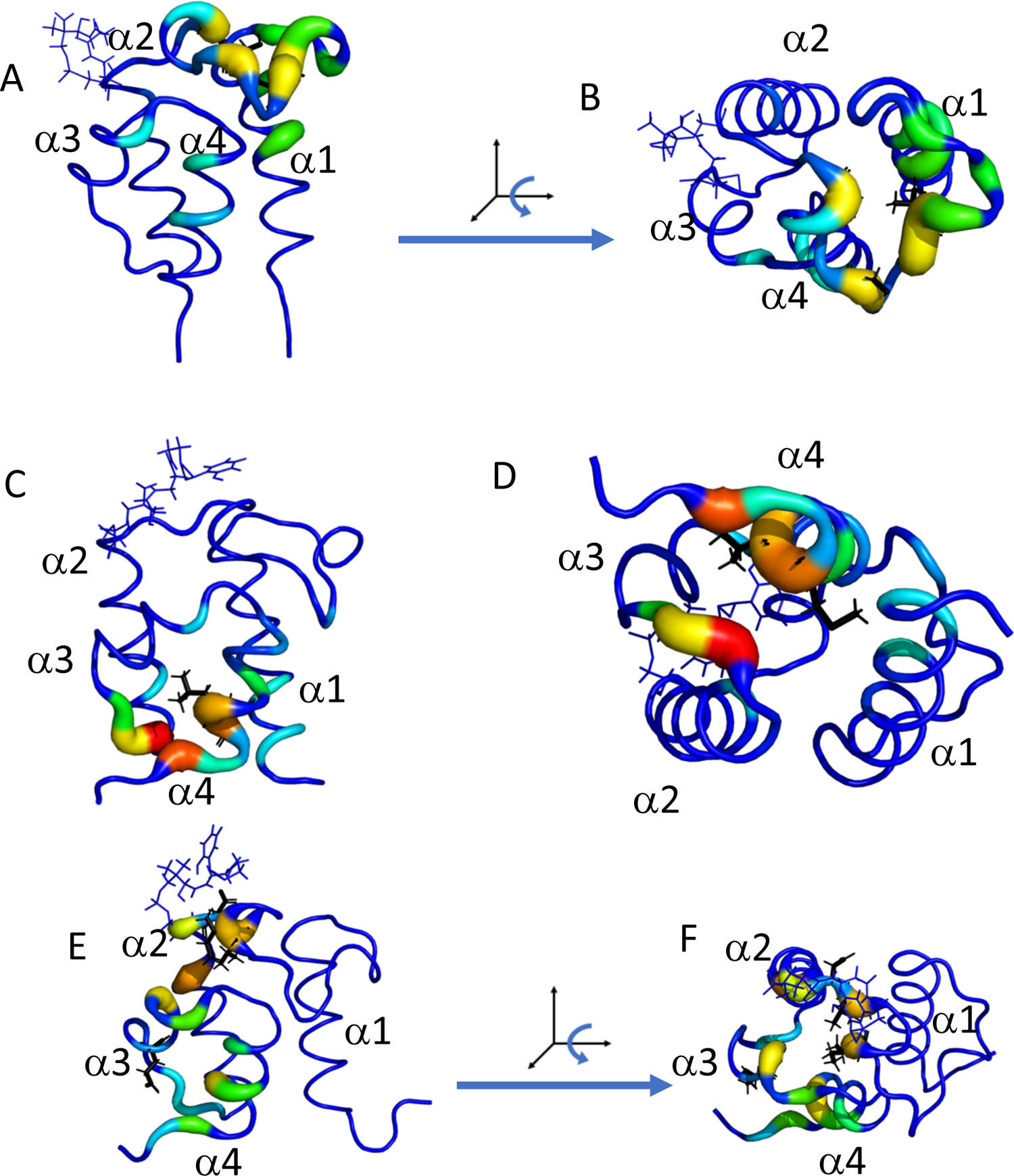
TACTICS predicts binding sites in the holo and loaded-ArCP. The PP arm is shown using blue sticks. A, B. TACTICS output for a predicted site in the holo-ArCP. C, D. TACTICS output showing a predicted site in the loaded-ArCP. E, F. TACTICS output showing another predicted site in the loaded-ArCP.

In the loaded-ArCP, TACTICS predicts a binding site that includes helix *α*4 and loop2 (between *α*2 and *α*3). This site is shown in Figure 6 C, D and Supplemental Figure S14. Potentially important residues on helix *α*4 include L86, M87, and R90. Potentially important residues on loop2 include Y67 and R68. This site provides an interesting contrast to the “Apo 2” site (seen in the apo and holo results). Helix *α*1 is a key part of the “Apo 2” site, but it plays a very minor role in the pocket shown in the loaded figure. The differences between the two pockets are caused by movement of the protein’s C terminus, as shown in Supplemental Figure S14 A.

TACTICS predicts another binding site for the loaded-ArCP. This site is shown in Figure 6 E, F and Supplemental Figure S15. The predicted site includes residues in loop1 (between *α*1 and *α*2), *α*2, and *α*3. Potentially important residues on loop1 and *α*2 include I46, L50, the S52-PP-salicylate arm, and L55. Y75 is a potentially important residue on *α*3.

## Conclusions

We created the TACTICS algorithm that analyzes MD simulations to find druggable sites in proteins. TACTICS first clusters the trajectory to select several frames for analysis. It then applies a random forest ML model to each selected frame. The model takes a protein conformation and scores each residue’s druggability in that conformation. The model was trained on a database of crystal structures; to capture conformational diversity, multiple structures were included for each protein. Sites identified by the ML algorithm undergo fragment docking to further characterize their bindability. To our knowledge, TACTICS is the only freely available program that both uses an ML model specifically designed for MD trajectories and accepts already-generated MD data. The TACTICS software is available at https://github.com/Albert-Lau-Lab/tactics_protein_analysis.

We applied the TACTICS algorithm to the SARS-CoV-2 main protease (nsp5) and 2’-O RNA methyltransferase (nsp10/16). In the main protease, TACTICS found the active site and multiple known allosteric sites; all of these sites have been observed experimentally. In the methyltransferase, TACTICS found the RNA and SAM binding sites and predicted the locations of sites not observed in experimental results.

We also applied TACTICS to the *Yersinia pestis* ArCP. The absence or presence of a large phosphopantetheine (PP) arm in apo or loaded and holo forms makes it a unique test case. TACTICS identifies several potential drug-binding sites that merit further study. Site predictions differ between the protein’s apo, loaded, and holo forms, suggesting possible differences between the forms’ bindability. The TACTICS results for this protein emphasize the software’s usefulness for proteins with unusual characteristics.

## Acknowledgements

We thank Dominique P. Frueh for helpful discussions. REST2 Simulations were carried out at the Argonne Leadership Computing Facility (ALCF), thanks to a grant from the COVID-19 High-Performance Computing Consortium (MCB200192). ArCP simulations were carried out at the Maryland Advanced Research Computing Center (MARCC) at Johns Hopkins University. This work was supported by the National Institutes of Health grant T32GM135131.

## Data and Software Availability

MD simulation data and TACTICS output is available from the Zenodo website; it can be found using the DOI 10.5281/zenodo.4538911. The TACTICS code is available at https://github.com/Albert-Lau-Lab/tactics_protein_analysis.

