## Supplemental Figures for "Finding Druggable Sites in Proteins using TACTICS"

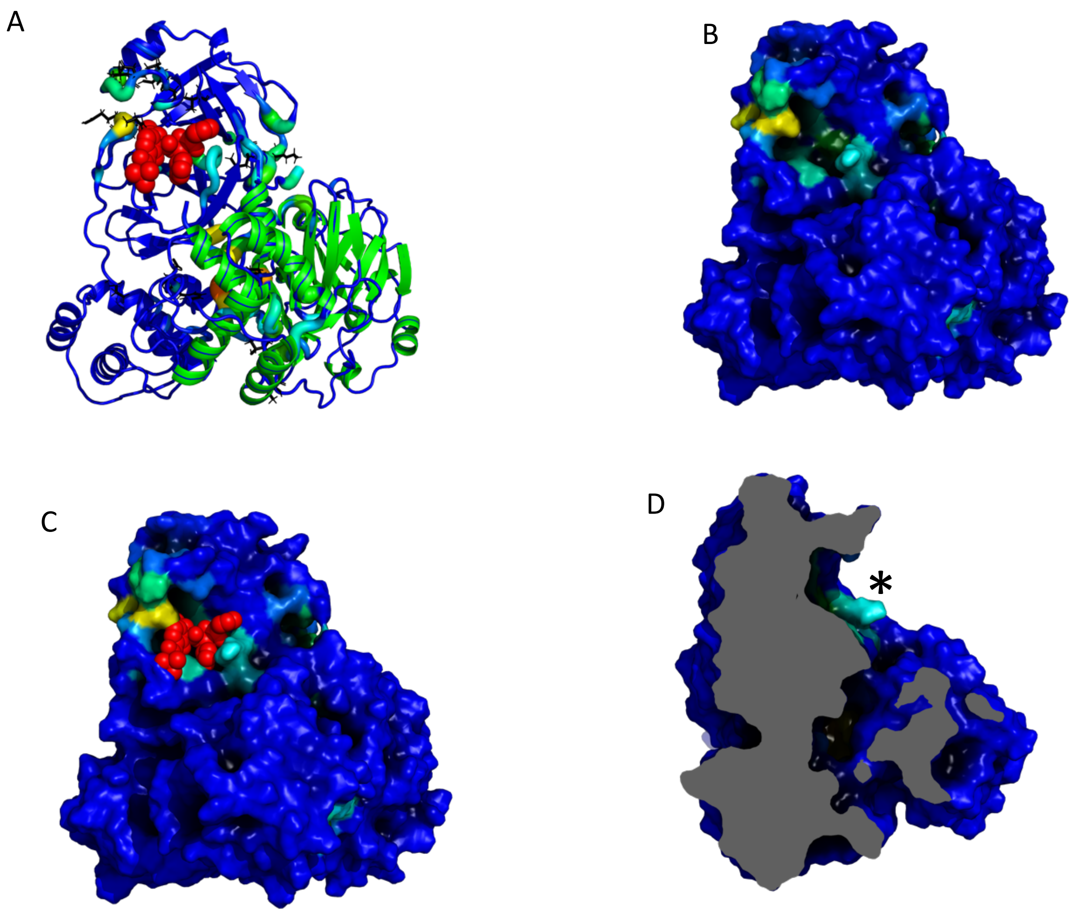


Supplemental Figure S1. Additional images of TACTICS output showing the SARS-CoV-2 main protease active site. A. The same as Figure (2B), with the protomers colored to show tertiary structure. B. Surface view of the pocket, with residues colored by TACTICS fragment docking scores. C. The same as (B), with an active-site inhibitor called 13b added from PDB ID 6Y2F. D. A rotated view of the protein. The asterisk indicates the active site location.

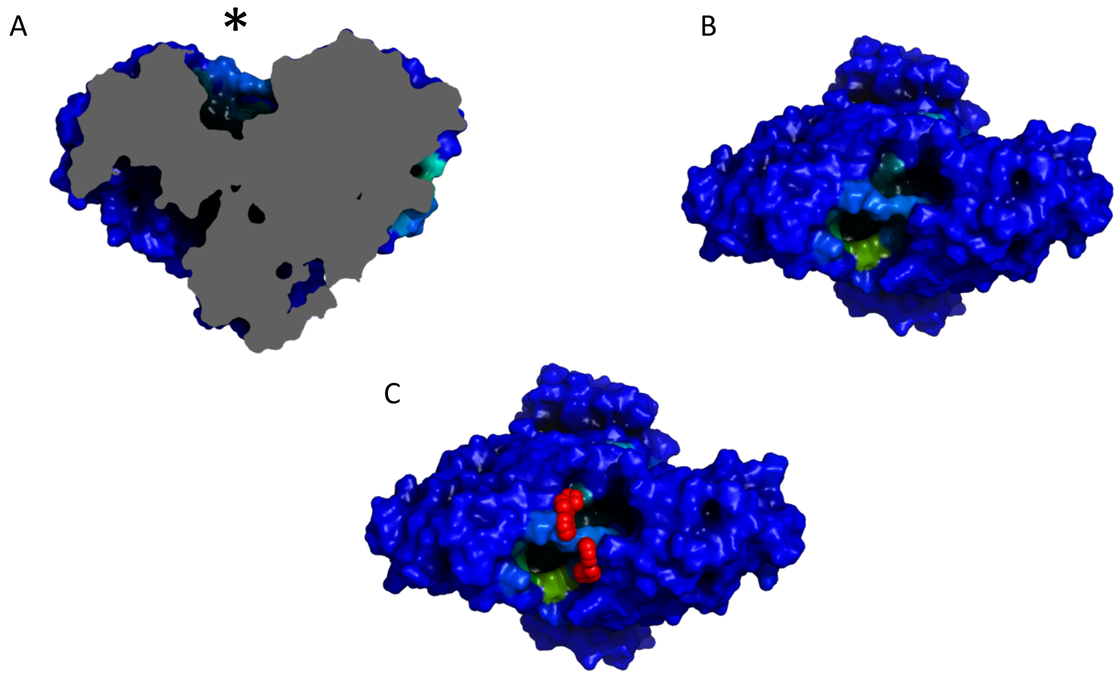


Supplemental Figure S2. Additional images of TACTICS output showing a known allosteric site in the SARS-CoV-2 main protease. A. A slice representation of the protein. The pocket location is indicated by the asterisk. B. Surface view of the top of the pocket, with residues colored by TACTICS fragment docking scores. C. The same as (B), with an inhibitor called x0425 or Z1401276297 added from PDB ID 5RGJ.

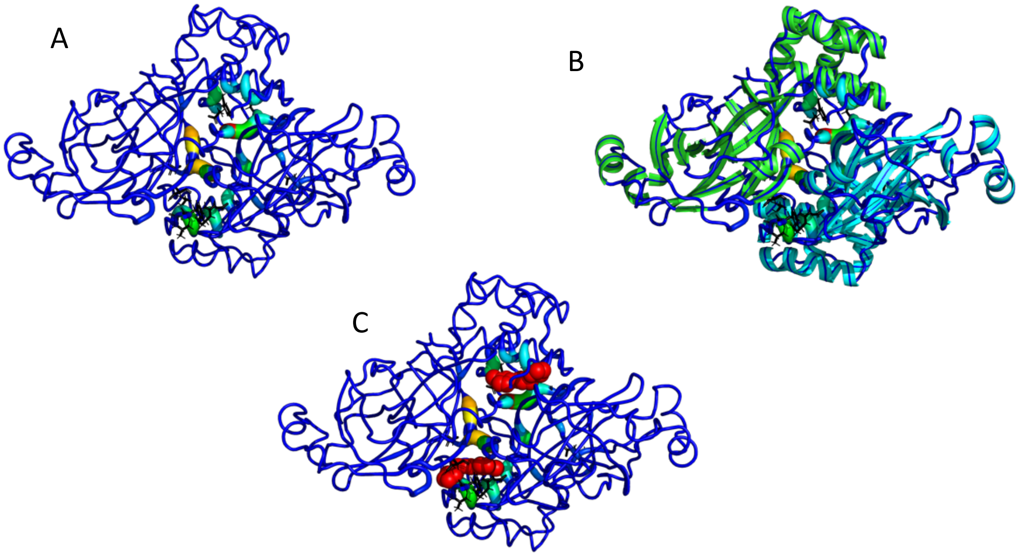


Supplemental Figure S3. TACTICS finds a known binding site at the SARS-CoV-2 main protease dimer interface. A. Top view of the TACTICS output. B. The same as (A), with the protomers colored to show the tertiary structure. C. The same as (A), with an inhibitor called x1187 or Z2643472210 added from PDB ID 5RFA.

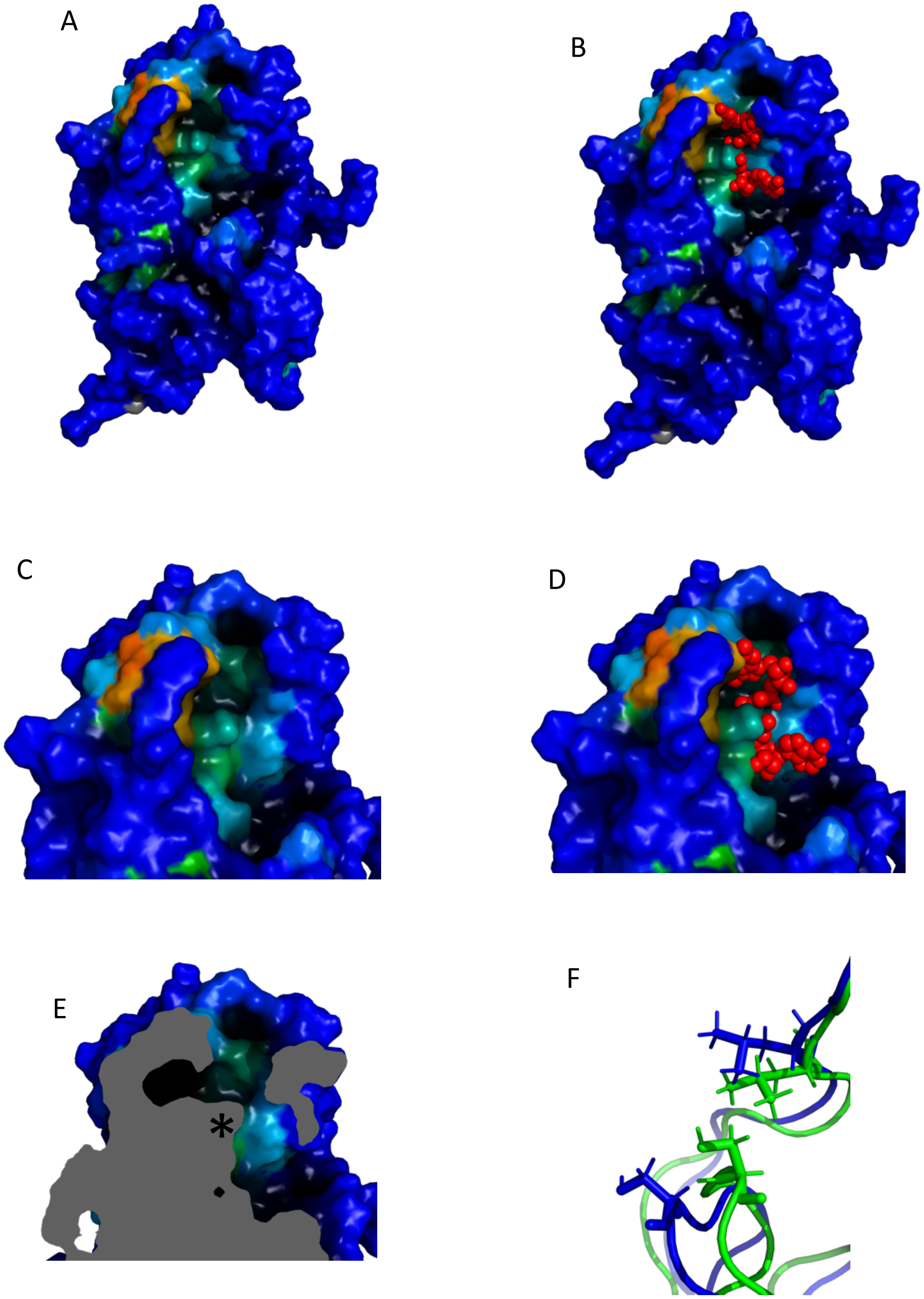


Supplemental Figure S4. Additional images of a predicted site connected to the MTase RNA binding site. A. Surface view of the structure shown in figure 3C. B. Surface view with m$^7$GpppA added from PDB structure 6WVN. C,D. Slightly rotated close-up of binding site. E. Slice view of site. The asterisk notes the area where the binding site is. Notice the cavity on the left; its opening is controlled by L27, S202, and nearby residues. F. An alignment of two frames from the MD data. L27 and S202 are shown in sticks. In the green frame, the residues are close to each other, blocking part of the cavity. In the blue frame, the residues are farther apart, allowing for access. Some parts of the protein were removed for image clarity.

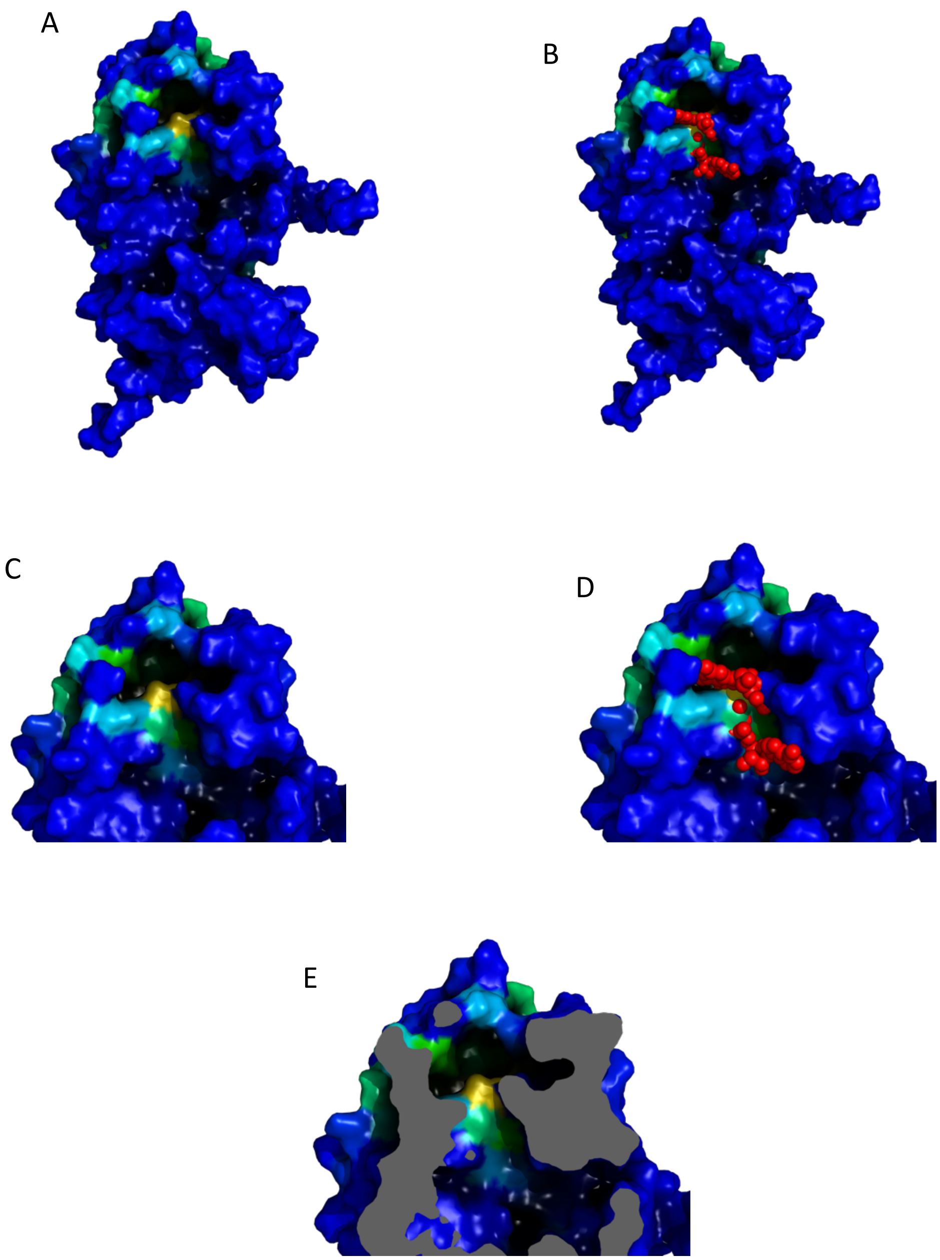


Supplemental Figure S5. Additional images of a predicted site connected to the MTase RNA binding site. This site is different from the one seen in Figure S4. A. Surface view of the structure shown in figure 3E. B. The same as (A), with $m^7$GpppA added from PDB structure 6WVN. C, D. Slightly rotated close-up of binding site. E. Slice view of site. Notice the cavity at the top right.

A B

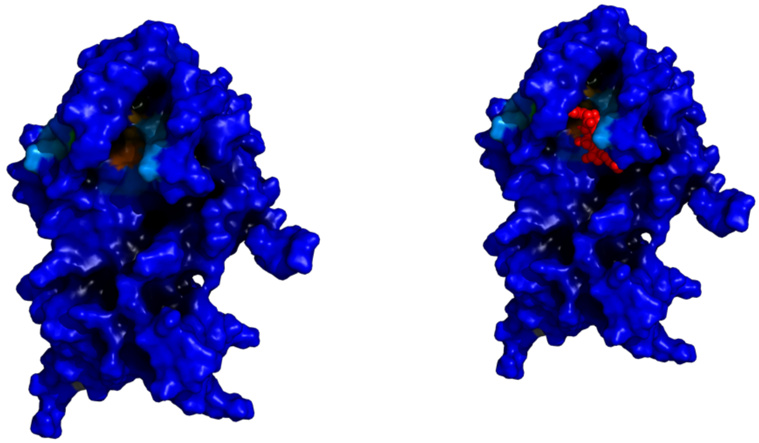

Supplemental Figure S6. Additional images of the SAM site as identified by TACTICS. A. A surface-representation of Figure 4A. B. A surface-representation version of Figure 4B.

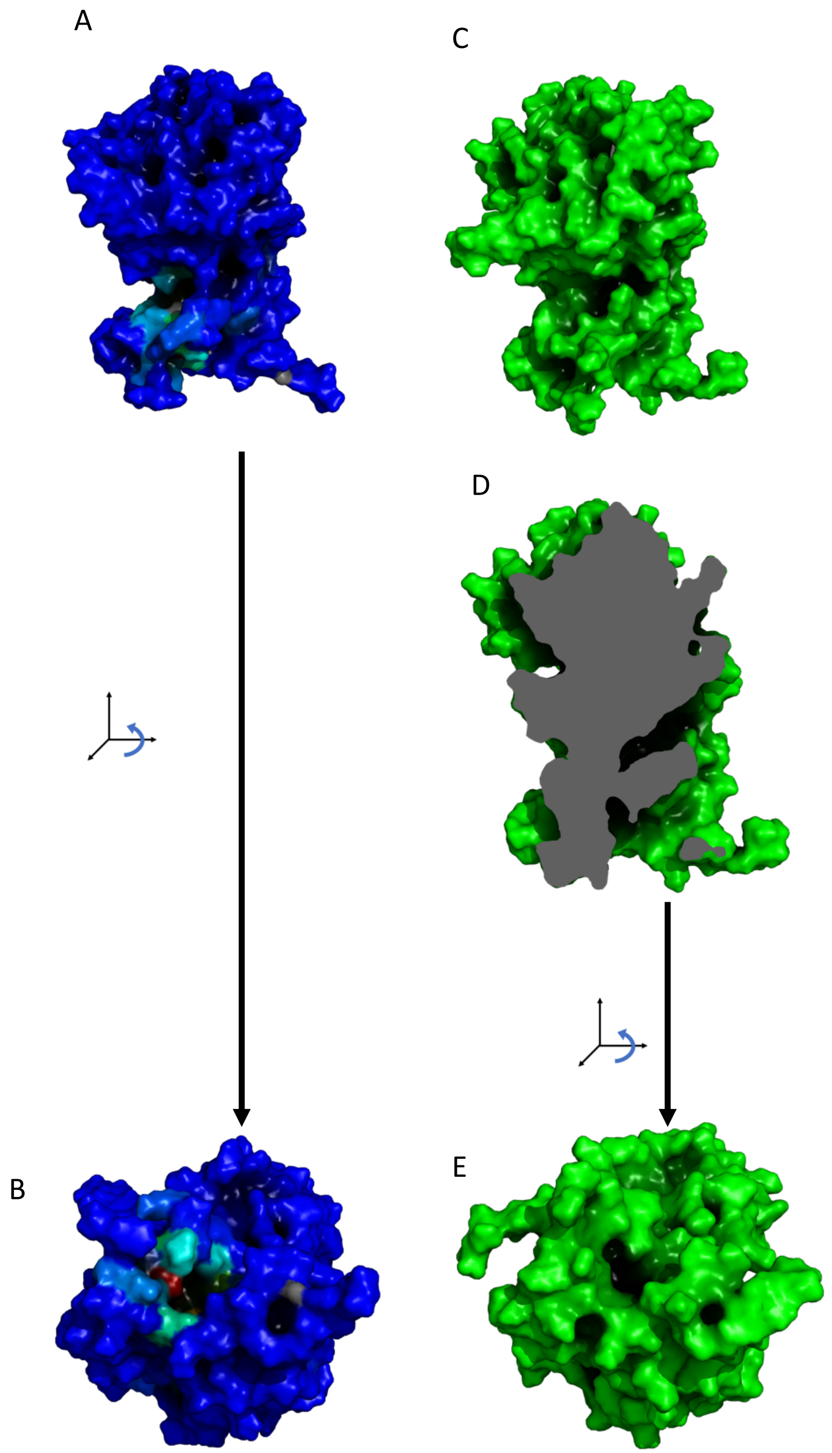


Supplemental Figure S7. Additional images of a predicted binding site at the dimer interface. A. A surface view of MTase. The predicted binding site is at the bottom left; it is colored by TACTICS. B. A rotated surface view showing the binding site. C. Surface view of the first frame of the MD data. D. Slice representation of the first frame of the MD data. Note that the tunnel is not fully present. E. Rotated view of the first frame.

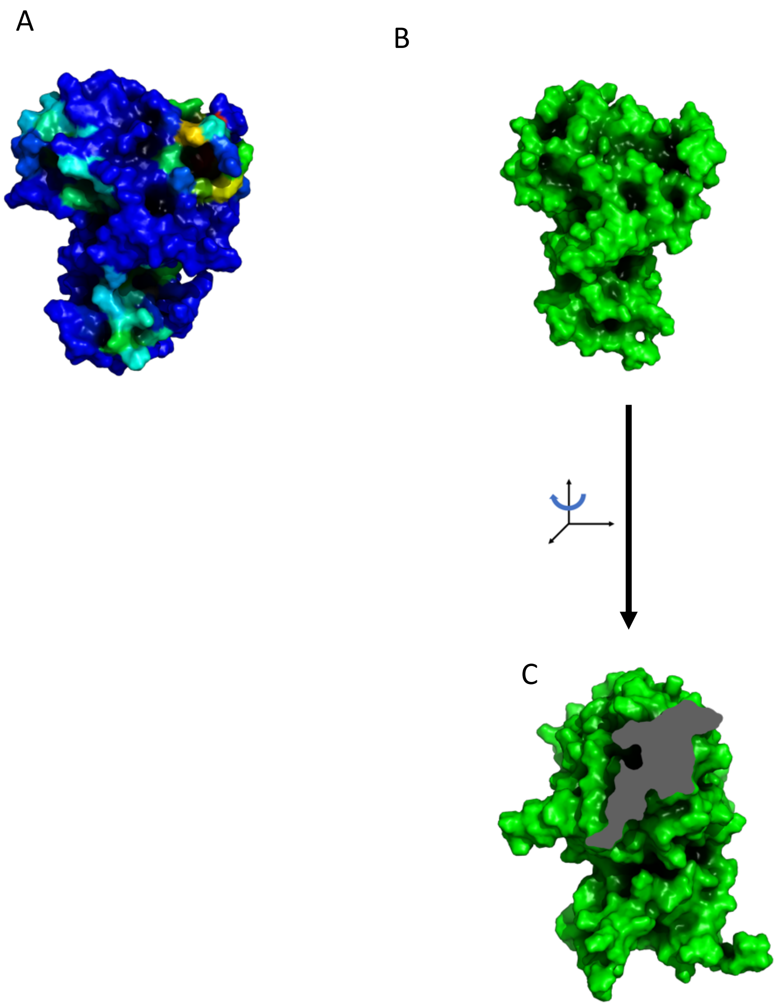


Supplemental Figure S8. Additional images of a predicted binding site on the methyltransferase. A. A surface view of MTase. The predicted site of interest here is on the right. The RNA site is on the left. B. Surface view of the first frame of the MD data. C. Slice representation of the first frame of the MD data.

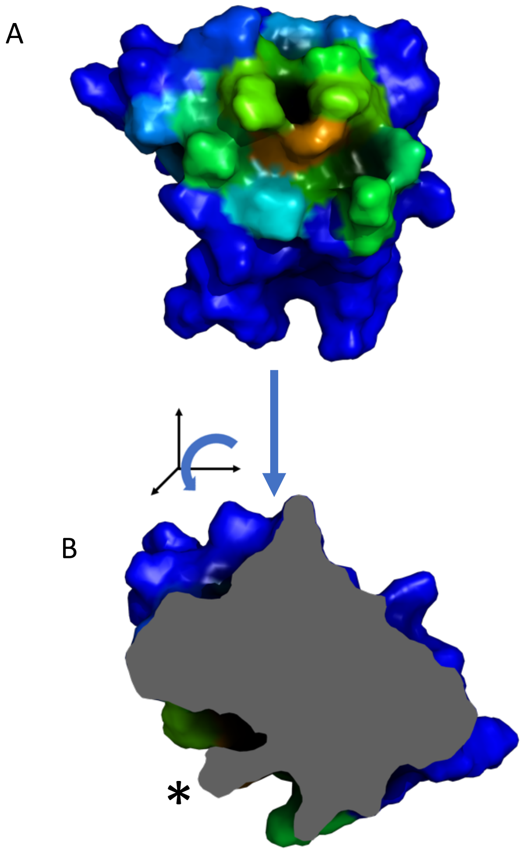


Supplementary Figure S9. Additional images of the apo-ArCP predicted site shown in Figure 5 A-B. A. A surface representation of Figure 5A. B. A rotated slice representation of this conformation. The asterisk marks the predicted site location. C. Surface view of the first frame of the MD trajectory, aligned with (A). D. Slice view of the first frame of the MD trajectory. The asterisk marks the predicted site location.

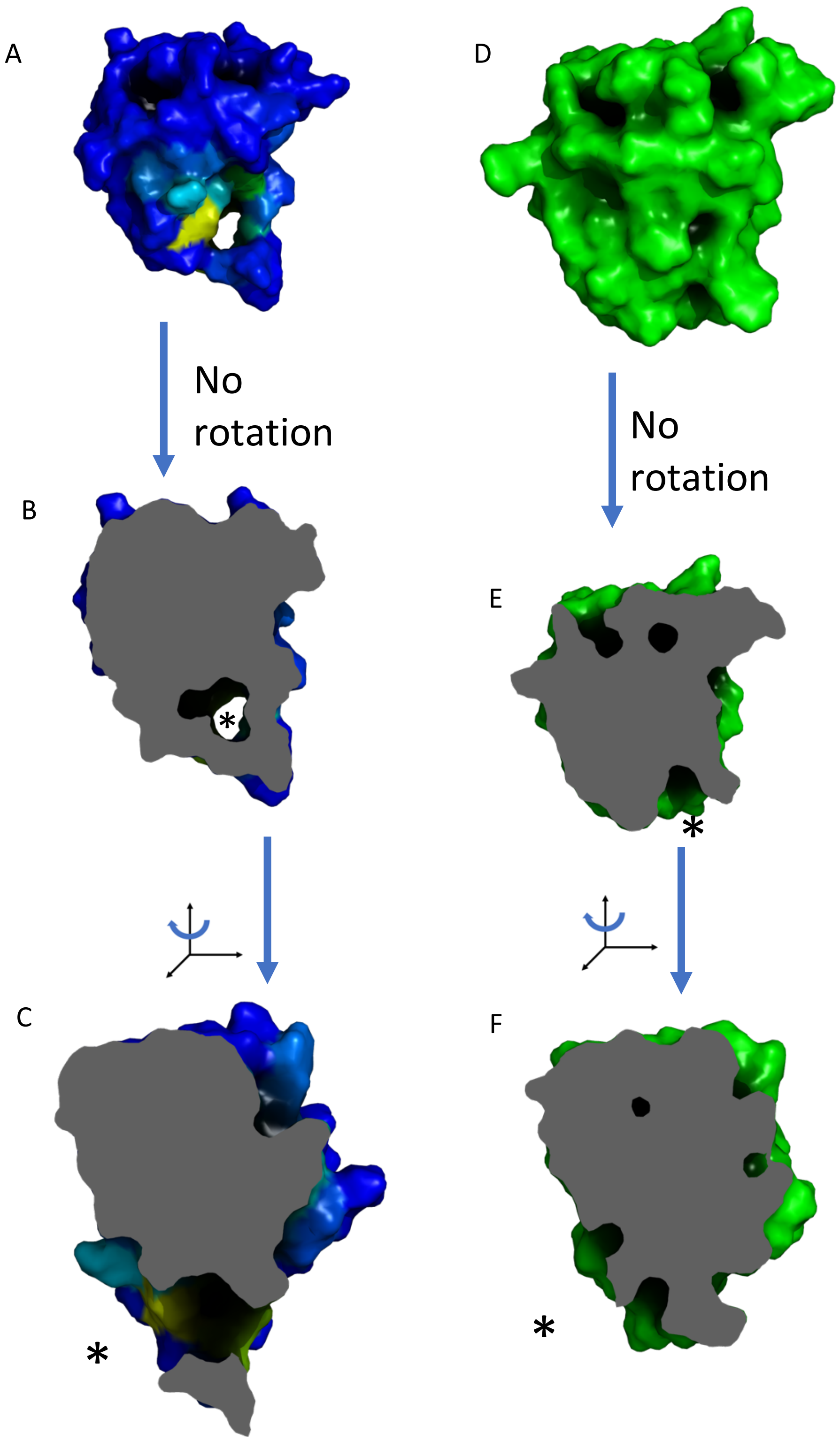


Supplementary Figure S10. Additional images of the apo-ArCP predicted site shown in Figure 5 C-D. A. A surface representation of Figure 5C. B, C. Slice views of this frame. The asterisk indicates the predicted site's location. D. Surface view of the first frame of the MD trajectory, aligned with (A). E, F. Slice views of the first frame. The asterisk indicates the predicted site's location.

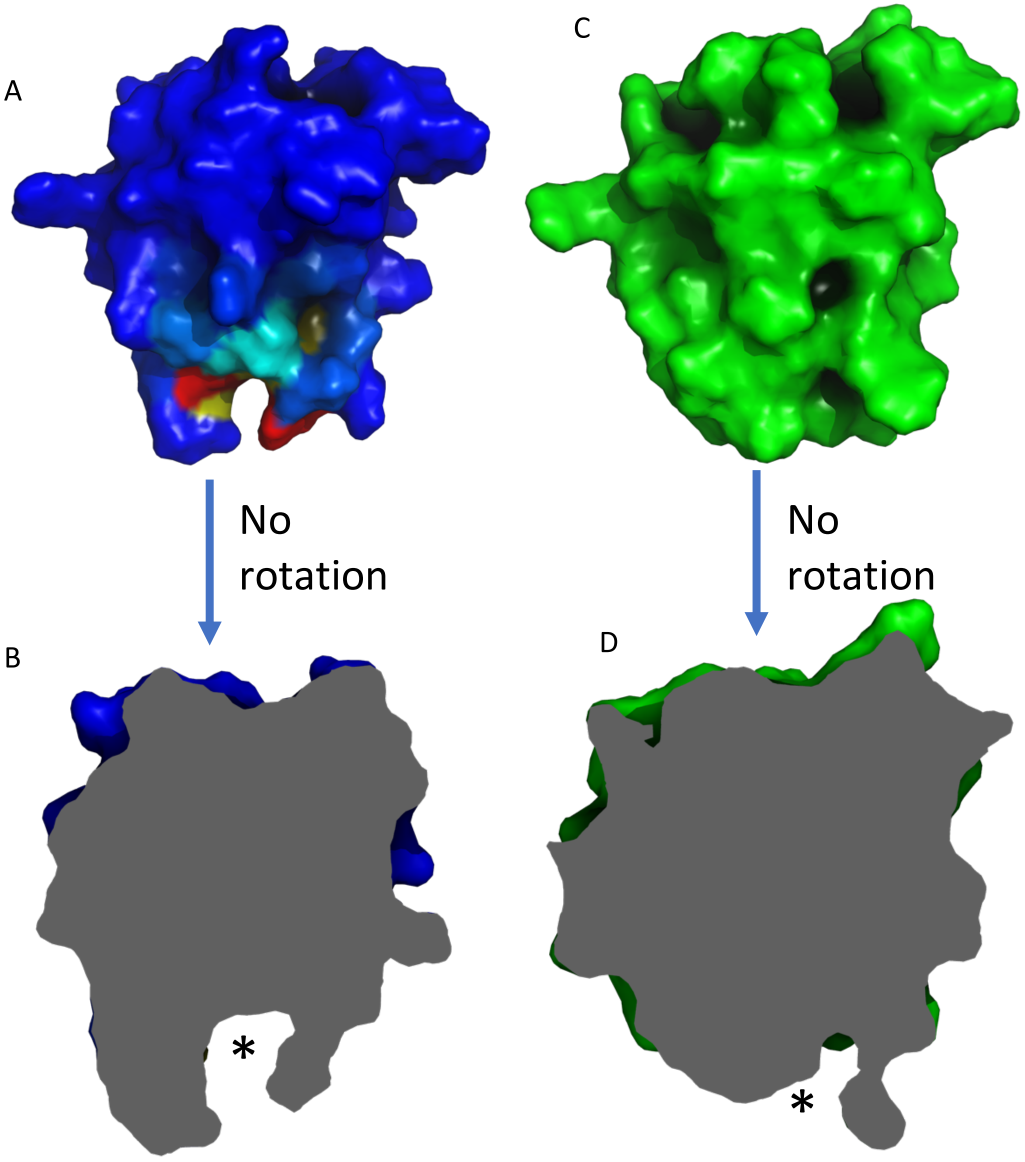


Supplementary Figure S11. Additional images of the apo-ArCP predicted site shown in Figure 5 E-F. A. A surface representation of Figure 5E. B. Slice views of this frame. The asterisk indicates the predicted site's location. C. Surface view of the first frame of the MD trajectory, aligned with (A). D. Slice view of the first frame. The asterisk indicates the predicted site's location.

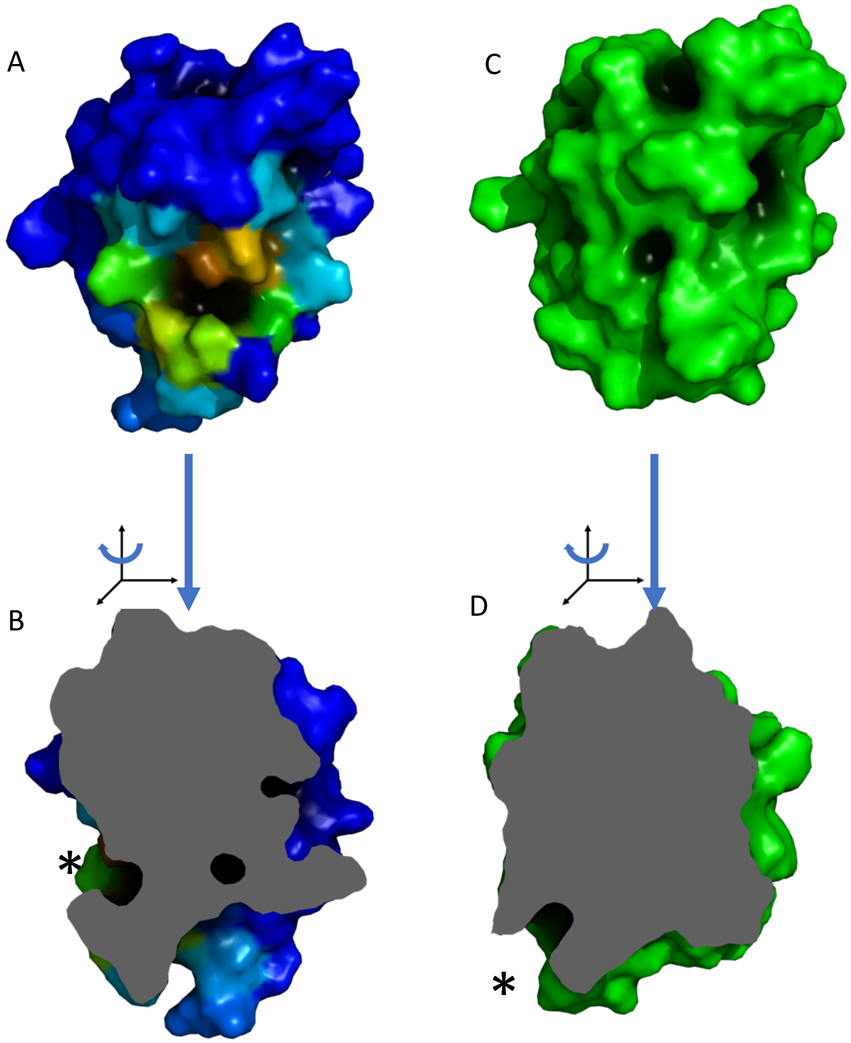


Supplementary Figure 12. Additional images of the apo-ArCP predicted site shown in Figure 5 G-H. A. A surface representation of Figure 5G. B. A rotated slice view of this conformation. The asterisk indicates the predicted site's location. C. Surface view of the first frame of the MD trajectory, aligned with (A). D. Slice view of the first frame. The asterisk indicates the predicted site's location.

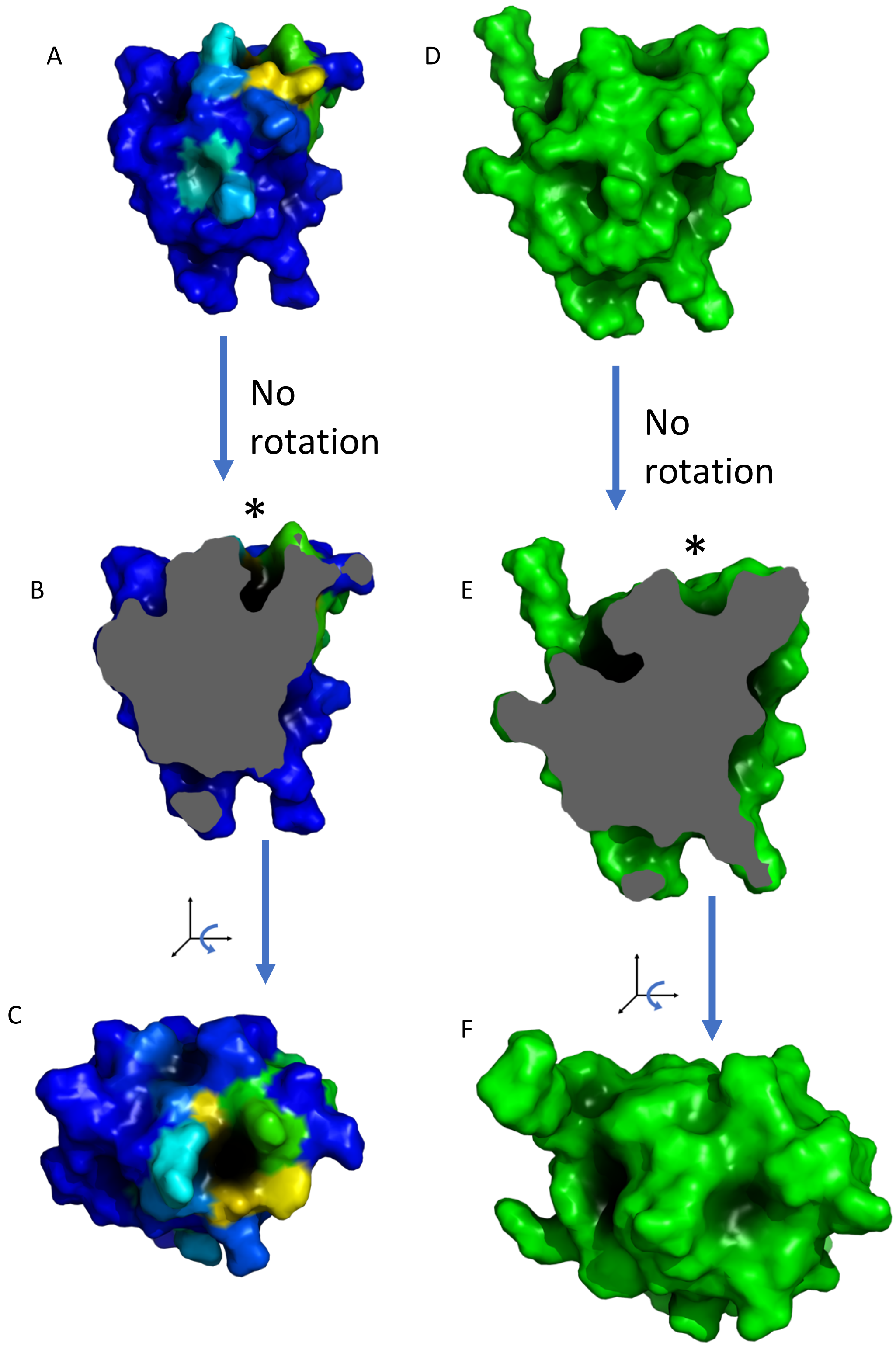


Supplementary Figure S13. Additional images of a predicted binding site in the holo-ArCP. A. Surface representation of Figure 6A. B. Slice representation of the site. The asterisk marks the site location. C. A rotated view of the site. D. Surface view of the first frame of the MD trajectory, aligned with (A). E. Slice view of the first frame. The asterisk indicates the predicted site's location. F. A rotated view of the first frame.

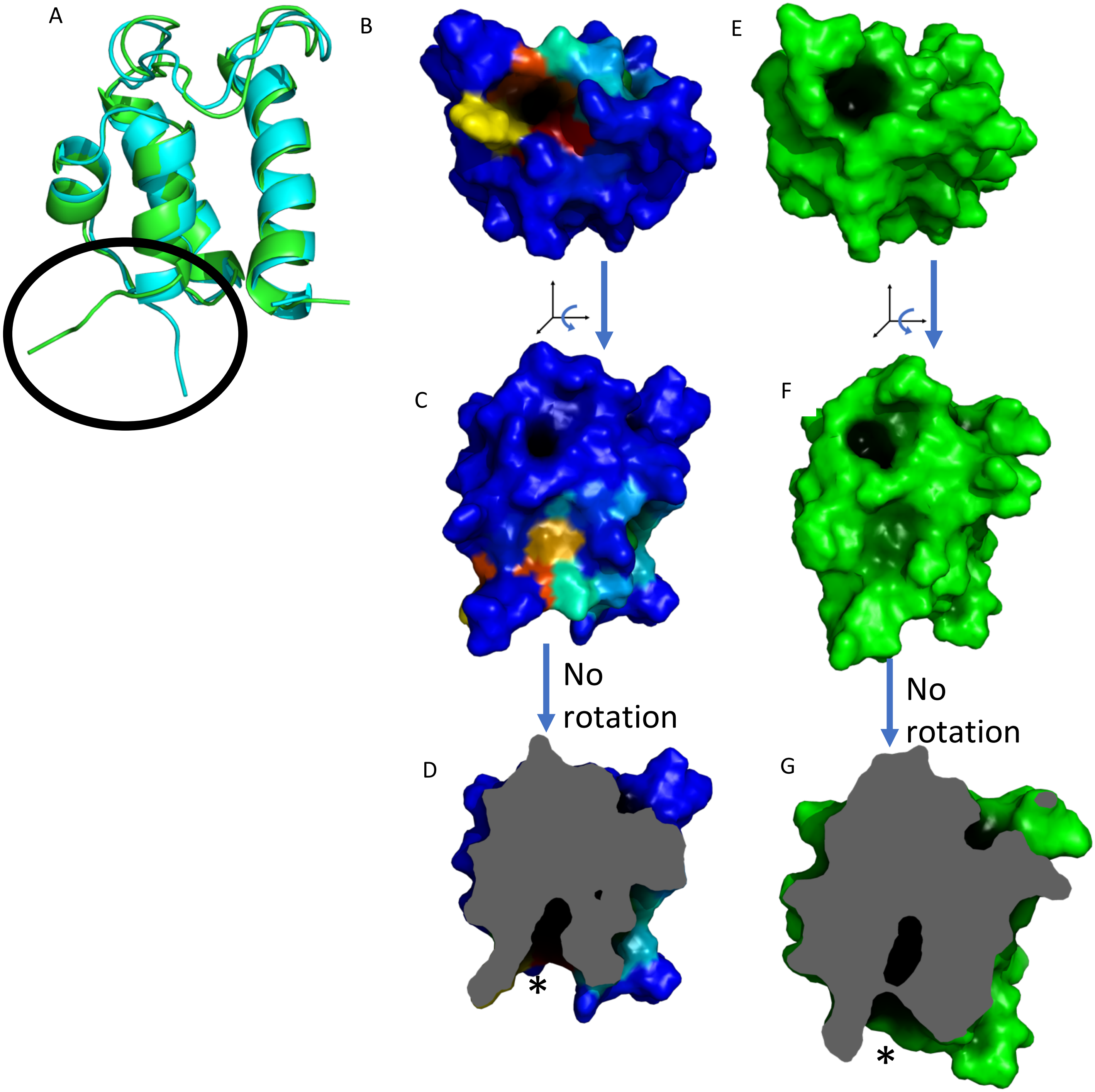


Supplementary Figure S14. Additional images of a predicted binding site in the loaded-ArCP. A. Alignment between a frame from the apo data (blue) and a frame from the loaded data (green). The C termini are circled to emphasize the dramatic difference in conformation. B. Surface representation of Figure 6D. C. Surface representation of Figure 6C. D. Slice view of the predicted site. The asterisk marks the site location. E. Surface view of the first frame of the MD trajectory, aligned with (B). F. Surface view of the first frame, aligned with (C). G. Slice view of the first frame. The asterisk marks the site location.

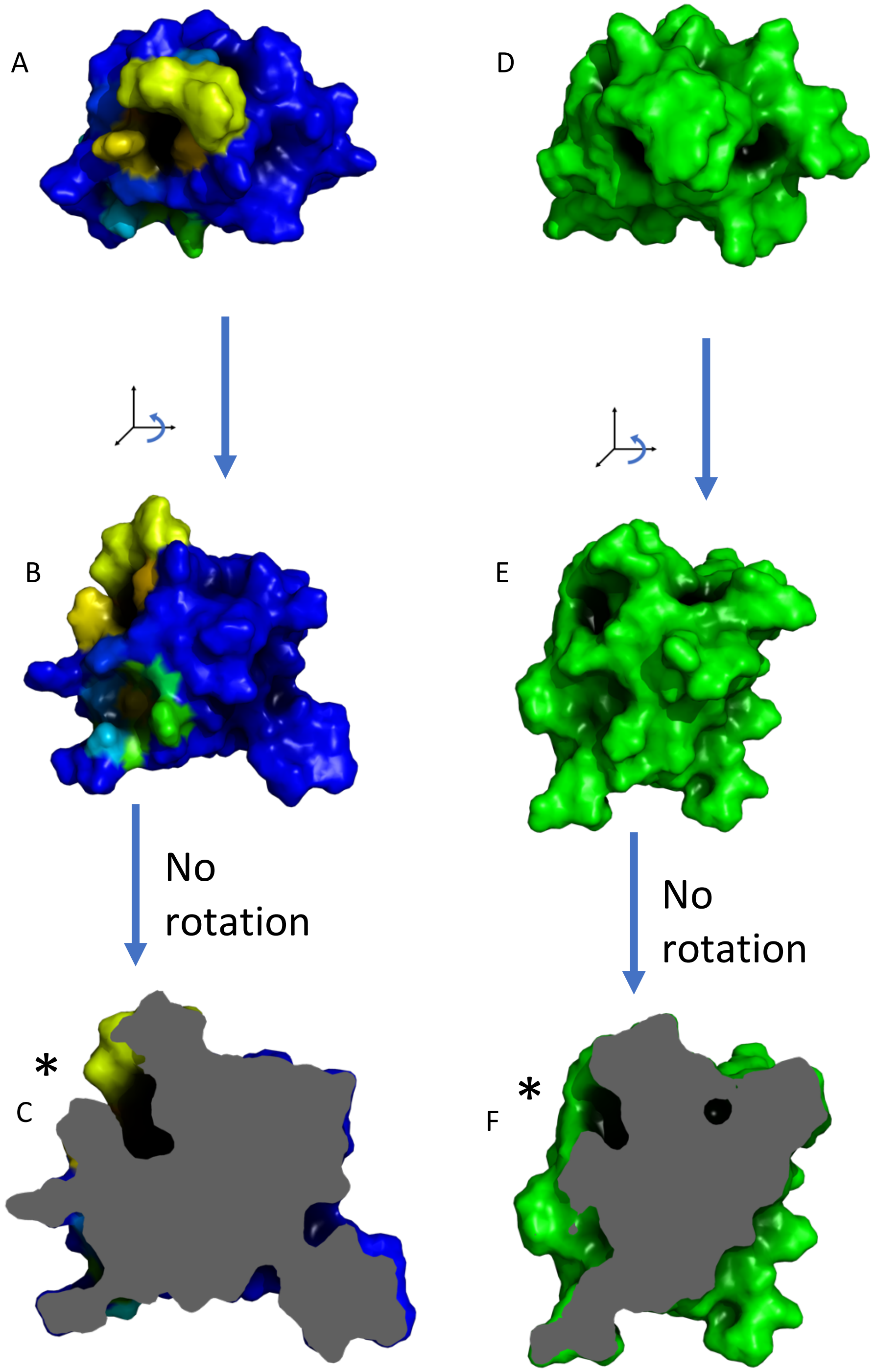


Supplementary Figure S15. Additional images of a predicted binding site in the loaded-ArCP. In the cartoon representations, the PP arm is shown using blue sticks. On the right, the green proteins are the initial structure (before MD simulation). A. Surface representation of Figure 6F. B. Surface representation of Figure 6E. C. Slice view of the predicted site. The asterisk marks the site location. D. Surface view of the first frame of the MD trajectory, aligned with (A). E. Surface view of the first frame, aligned with (B). F. Slice view of the first frame. The asterisk marks the site location.
